# Microevolution of clade II isolates of *Candida auris* highlights multifaceted intra-clade heterogeneity in acquiring resistance towards amphotericin B

**DOI:** 10.1101/2023.09.20.558730

**Authors:** Anshu Chauhan, Praveen Kumar, Mohit Kumar, Aswathy Narayanan, Kusum Yadav, Basharat Ali, Amandeep Saini, Ashutosh Singh, Atanu Banerjee, Shivaprakash M. Rudramurthy, Arunaloke Chakrabarti, Alok K. Mondal, Naseem A. Gaur, Kaustuv Sanyal, Rajendra Prasad

## Abstract

*Candida auris* exhibits high-level resistance to amphotericin B (AmB). Mechanisms such as ergosterol biosynthesis malfunction, oxidative damage mismanagement, and increased drug efflux contribute to AmB resistance in *C. auris*. In this study, we experimentally evolved two East Asian drug-susceptible clade II isolates of *C. auris* (P2428 and CBS10913^T^) isolated from different geographical locations to develop resistance against AmB. We analysed alterations in karyotype, genome sequence, and gene expression profiles to uncover the mechanisms driving AmB resistance. The independently evolved clade II adaptors displayed up to 4-16-fold higher MIC^50^, as compared to the parental cells. *AOX2* (alternative oxidase) and the cell wall integrity pathway have been identified as critical in the development of AmB resistance. However, we noted certain intra-clade heterogeneity in the associated mechanisms. While in P2428 adaptors (P-lines), the ergosterol and sphingolipid pathways appear to play a crucial role, this was not the case for CBS10913^T^ adaptors (A-lines), which acquired resistance independent of lipid-associated changes. The transcriptomic, WGS, and phenotypic analyses also confirm that the evolved AmB-resistant isolates follow distinct trajectories for adaptation, Furthermore, unlike the fluconazole-resistant isolates, as reported previously, changes in ploidy do not seem to contribute to the differential mechanisms of AmB resistance. Overall, this study not only provides insights into the mechanisms and pathways involved in AmB resistance but also highlights intra-clade-heterogeneity that exists within clade II of *C. auris*.

**Importance:** *Candida auris* demonstrates significant resistance to amphotericin B (AmB) that stems from factors like alteration of ergosterol biosynthesis, perturbation of the oxidative damage response, etc. A comprehensive understanding of underlying mechanisms can be studied in a holistic manner by subjecting resistant as well as susceptible clinical isolates to a comparative genome-level analysis. An alternate and more dynamic approach is to expose susceptible isolates to a certain concentration of drug which is not lethal but can trigger the resistance mechanisms. In the present study, we evolved *C. auris* towards AmB and observed novel and differential mechanisms of resistance development, in two different isolates despite belonging to the same clade. This study provides insights into the intra-clade heterogeneous behavior of *C. auris* towards AmB.

## Introduction

The emergence of multidrug-resistant *Candida auris* across multiple continents is a global concern. The presence of different clonal isolates suggests the independent evolution of drug resistance in response to antifungal therapies*. C. auris* poses unique challenges compared to other pathogenic *Candida* species, particularly due to its resistance to multiple classes of antifungal agents (1, 2). The *C. auris* isolates typically show uniform resistance to fluconazole, approximately 50% resistance to voriconazole, around 30% resistance to amphotericin B (AmB), and nearly 10% resistance to echinocandins (3). However, in the fight against *Candida* infections, four major classes of antifungal drugs, namely azoles, polyenes, nucleic acid analogs, and echinocandins, remain the frontline treatments (4, 5). Azoles, which target the enzyme Erg11p involved in the production of ergosterol, are the most used antifungals. They cause changes in the fungal cell membrane by reducing ergosterol levels and increasing the presence of toxic sterol precursors. Echinocandins, a relatively new class of antifungals, target the fungal cell wall by inhibiting the synthesis of 1,3-β-D-glucan, which is encoded by FKS genes. Mutations in FKS1/2 genes are the main cause of echinocandin ineffectiveness in *Candida* cells (6). Additionally, compounds that inhibit nucleic acids like flucytosine also exhibit antifungal activity (7).

Polyenes, such as AmB and nystatin (NYT), are commonly used when azole therapy fails. They target ergosterol and genes involved in its biosynthesis (4). The mechanism of action of AmB is explained by the sterol sponge model, where the drug removes ergosterol from lipid bilayers, leading to its accumulation outside the membrane (8). The action of AmB, stands apart from the conventional mode of drug action, as it functions through the unique mechanisms of ergosterol sequestration and induction of oxidative stress (9). Unlike many other antifungal drugs targeting specific enzymes, AmB exerts its effects by binding to ergosterol, a crucial component of the fungal cell membrane, forming pores and compromising the integrity of the membrane. This disruption leads to the leakage of cellular contents and, importantly, triggers oxidative stress within the fungal cell. Curiously, AmB resistance in *C. auris* and other *Candida* species remains one of the most enigmatic and least comprehended drug resistance mechanisms. This is primarily due to the distinctive nature of AmB action, which contrasts with the conventional enzyme inhibition seen in other drug resistance scenarios. The complexity of the underlying mechanisms has contributed to the challenges in unravelling the precise factors that drive resistance. Up to this point, only a handful of insights have emerged concerning the intricate landscape of AmB resistance in *C. auris*. Notably, increased expression of specific genes involved in ergosterol biosynthesis, namely *ERG1*, *ERG2*, and *ERG6*, has been observed in the context of resistance. This heightened expression suggests a potential compensatory response to counter the disruption caused by AmB’s interference with ergosterol, the cell membrane’s vital component (9–11).

Natamycin, another polyene, has been found to interact with membrane-localized amino acids and sugar transporters, as well as it affects the ergosterol-dependent membrane permeabilization (12). However, the mechanisms of action and development of resistance against polyenes, including AmB, are still not fully understood. Clinical isolates of *C. auris* have shown increased resistance to AmB, possibly due to malfunctioning in ergosterol biosynthesis, faulty management of oxidative damage, and elevated efflux (13). TOR kinases and ROS levels have also been linked to AmB susceptibility (14). Studies have also pointed towards the DNA damage checkpoint proteins like *MEC3* to be involved in developing AmB resistance even though this remains to be confirmed (15–17). Further, identifying single nucleotide polymorphisms (SNPs) within essential genes has provided additional clues. Specifically, SNPs in *ERG2*, which encodes an enzyme involved in ergosterol biosynthesis, and *FLO8*, which is a transcription factor known to positively regulate the *ERG11* expression (18) and have been associated with AmB resistance. However, the current understanding of resistance mechanisms does not fully explain the high occurrence of AmB resistance in *C. auris* clinical isolates.

In this study, two susceptible East Asian isolates of *C. auris* of clade II, isolated from two different geographical locations, were adapted to higher concentrations of AmB by using the experimental evolution approach. While P2428 was recovered from pus of a diabetic patient in India, CBS10913^T^ was recovered in Japan from the external ear canal of an inpatient (19). The cells were exposed to a constant sub-lethal concentration of AmB for up to 100 generations. The adapted cells from both the progenitors showed up to 32-folds higher minimum inhibitory concentration (MIC_50_) compared to their original forms. Importantly, the adapted cells maintained their higher MICs even after several passages in drug-free media. The data suggests that two different drug susceptible isolates of clade II cells employ different transcription trajectories to develop high resistance to AmB exhibiting intra-clade heterogeneity.

## Results

### Selection of two *C. auris* susceptible isolates of East-Asian clade for microevolutionary studies towards Amphotericin B

P2428 was isolated from a diabetic patient’s pus in India and was initially expected to be a clade I isolate owing to the area of its isolation. Strikingly, analysis of P2428 using clade-specific primers (20) yielded amplicons only with clade-II primers, suggesting that P2428 is a clade II isolate. The raw reads from Illumina sequencing of P2428 were then mapped to the GenBank assemblies GCA_002759435.1 (clade I, strain B11205), GCA_003013715.1 (clade II, strain B11220), GCA_016772215.1 (clade III, strain B12037), GCA_008275145.1 (clade IV, strain B11245), and GCA_016809505.1 (clade V, strain IFRC2087). The variant calling was performed using the tool Snippy at Galaxy (usegalaxy.org) using default settings. 60606 single nucleotide variations (SNVs) were present between P2428 and the clade I strain, 59423 SNVs between P-2428 and the clade III strain, 157795 SNVs between P2428 and the clade IV strain, and 2346699 SNVs between P-2428 and the clade V strain. P2428 harboured 3701 SNVs compared to the clade II strain. This analysis cemented our observation that P-2428, isolated in India, belongs to clade II, contrary to the expectation. Another strain, CBS10913^T^ was also assessed for the clade identification (Fig. S1B). Its already reported in (21) that CBS10913^T^ is closely similar to the clade II reference strain B11220 (Fig. S1). This is the first report, to the best of our knowledge, that suggests *C. auris* clades other than Clade I are in circulation in the Indian subcontinent.

### Microevolution reveals the evolvability of clade II isolates of *C. auris* to acquire amphotericin B resistance

The two drug susceptible clade II isolates of *C. auris,* P2428 having MIC_50_ value 16 µgmL^-1^ for FLC and 0.5-1 µgmL^-1^ for AmB, and CBS10913^T^ with MIC_50_ value 8 µgmL^-1^ for FLC and 0.5 µgmL^-1^ for AmB were then subjected to experimental evolution by constantly exposing each of them to AmB concentration equivalent to their MIC_50_ values for 100 generations (Fig. S2A) (for details, see methods). All the evolved cell lines thus obtained after completion of 100 generations were harvested and subjected to AmB susceptibility testing through microdilution assays. These adapted replicates were designated as P-lines (evolved from P2428) and A-lines (evolved from CBS10913^T^). Two of the three P-line adapted strains (PA1 and PA2) exhibited a 16-32-fold increase in MIC_50_ values and one replicate (PA3) achieved up to 64-fold increased MIC_50_ value as compared to the control replicates. Whereas in case of the A-line adaptors (A1, A2, and A3), exhibited only 4-8-fold increase in MIC_50_ values as compared to the control replicates.

Following this, we assessed if the adapted replicates i.e., PA1, PA2, and PA3 belonging to P-line and A1, A2, and A3 belonging to A-line exhibiting high MICs towards AmB could be retained over a long period of time. This was tested by passaging the adapted replicates for 30 days on YEPD media plates at 24 h, in the absence of the drug, along with the parallelly run control replicates. Following the 30 days of drug-free daily passaging, all the P-line adaptors majorly retained the high MICs achieved after microevolution (Fig. S2 B). While, the A-line adaptors exhibited an ∼50% drop in AmB MIC_50_ (8 µgmL^-1^ to 2-4 µgmL^-1^) (Fig. S2 C). It was apparent that the isolates of the same clade not only responded differently to the experimental evolution regimen but their subsequent behaviour also differed. For further analysis, 3 random colonies from each set of passaged adaptors, displaying highest MIC_50_ values were selected and in A-line adaptors all the colonies of A3 replicates had exhibited drop in MIC values post drug free passages, hence 3 colonies were selected from A1, and A2 only (highlighted in different colours in the tables (Fig. S2 B, and C). These finally selected colonies were subjected to susceptibility testing by broth microdilution, and spot assays and found to be resistant towards AmB for up to 32 µgmL^-1^ in P-line and 8 µgmL^-1^ in the A-line adaptors (Fig. 1A, B, and 1 E, F). These resistant adaptors of A and P-lines were also tested for other polyene compounds, nystatin and natamycin and they were found to be cross-resistant towards these as well (Fig. S3), suggesting that the resistance developed is against the polyene class of drug in general.

**FIG 1:**
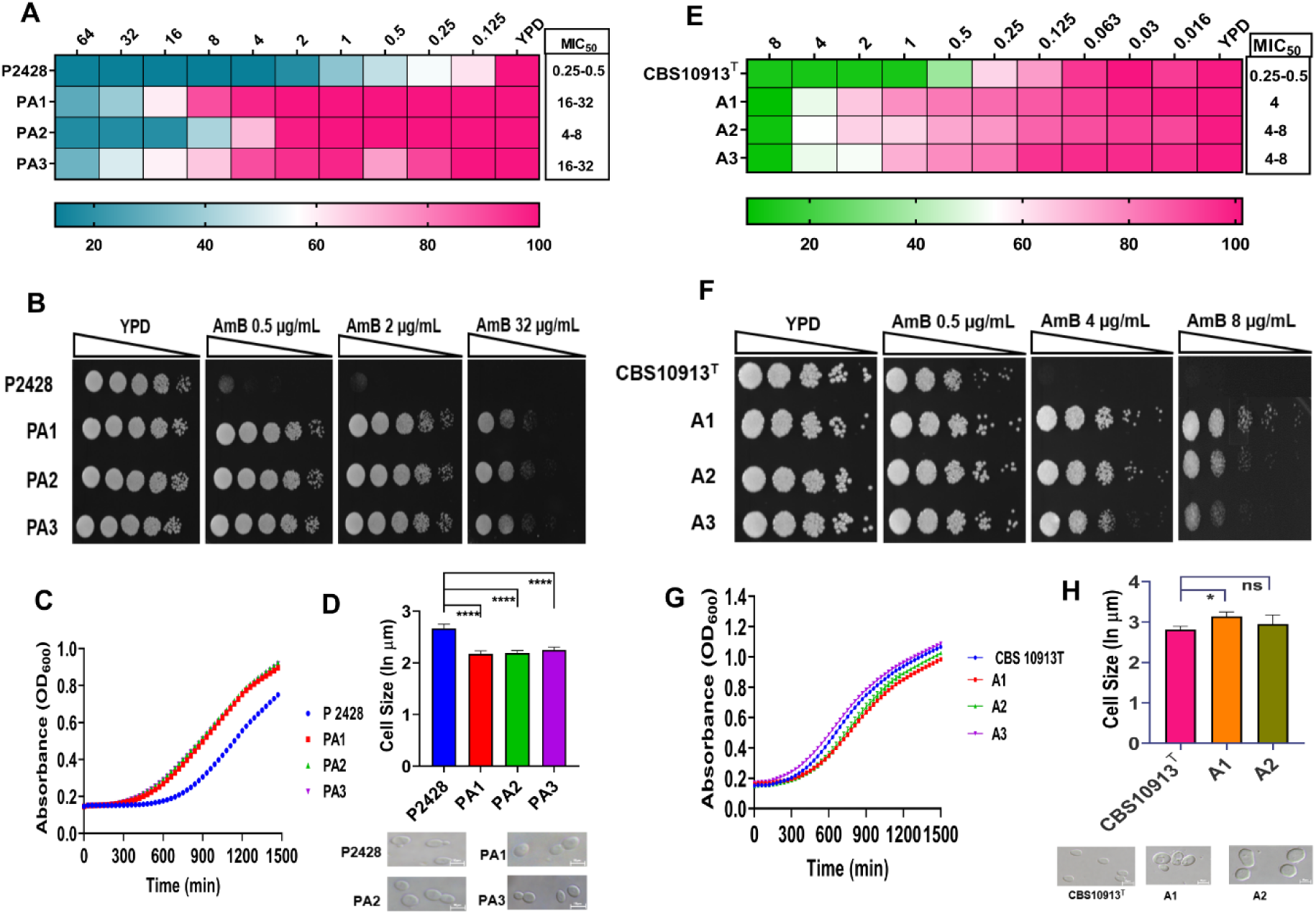
Directed evolution regime reveals evolvability of two different *C. auris* strains belonging to clade II towards AmB resistance. **(A, E)** Selected colonies exhibiting high resistance were checked by broth microdilution assay, results were recorded in a BioRad iMark microplate reader. **(B, F)** spot assays on YEPD agar plates with (0.5 µgmL^-1^, 4 µgmL^-1^, 8 µgmL^-1^,16 µgmL^-1^, and 32 µgmL^-1^ AmB) and without AmB. Growth differences were recorded after 48 hrs of incubation at 30°C by BioRad XR+ Gel documentation system. **(C, G)** Growth kinetics of P and A-lines adaptors were determined by a micro-cultivation method in a 96-well round bottom plate using a multimode microplate reader (Tecan Infinite M Plex, USA) in YEPD broth at 30 °C. The experiment was performed in triplicates and the mean values were plotted. **(D, H)** Differences in the cell sizes of the adaptors, compared to their respective parental strains, were calculated by using exponential growth phase cells in a microscope (Nikon A1 R fluorescence microscope, Japan). The cell sizes were calculated by taking an average of 100 cells counted in the NIS software in which the frames were kept uniform with a scale of 10 µm.

### AmB-resistant A-line evolved strains display an increase in cell size while both adapted lines do not show a change in growth

Unlike previous studies, this acquired resistance towards AmB appeared to be independent of growth defects.(Fig.1 C, and G). All the adapted replicates of A- and P-lines were checked for their growth rates and cell size. A-line adaptors showed an increase in cell size as compared to the parallelly run control strains (Fig. 1H), while the P-line adaptors showed a reduction in cell size (Fig. 1D). The P-line adaptors exhibited increased growth than the control (P2428), implying an increased fitness. However, A-line adaptors showed no significant difference in growth.

### AmB resistant A- and P-line adaptors display no collateral-resistance to azoles

The selected three colonies from each progenitor were evaluated for any cross-resistance towards other classes of antifungal agents. Among the azoles, fluconazole, voriconazole, clotrimazole, ketoconazole, and itraconazole, were tested. The AmB-adapted colonies of both the progenitors showed no cross-resistance towards the tested azoles (Fig. S3). Notably, the AmB-resistant adaptors of both the P, and A lines exhibited an 8-16-fold increase in MIC_50_ values towards other polyenes, nystatin and natamycin.

### Global Transcriptomic analysis of AmB-resistant adaptors show variable number of DEGs

To understand the molecular basis of the increased resistance to AmB in the evolved P and A lines, we performed global transcriptomic profiling. For this, the total RNA was extracted from all the three adaptors of each line by harvesting cells growing in an exponential phase in the absence of AmB. Each replicate was compared with the AmB-susceptible *C. auris* control, which was parallelly grown for 100 generations in the absence of the AmB. We used a 1.5-log_2_ fold change in the expression as the threshold for differentially expressed genes with an associated *p-value* of ≤0.05 as significant for our analysis. The global transcriptome of P-line and A-line adaptors revealed that the number of differentially expressed genes (DEGs) were variable among the replicates. Despite the variation in the number of differentially expressed genes, there was an appreciable similarity of genes among the three replicates of each progenitor. Likewise, a set of common 26 up-regulated and 25 down-regulated genes in P-line adaptors (Fig. 2 B, C), and 267 up- and 145 down-regulated genes were found in all the three adaptors of A-line (Fig. 2 E, F). This information of genes that are differentially regulated but are common in all three adaptors of each progenitor is depicted in (Fig. S4).

**FIG 2:**
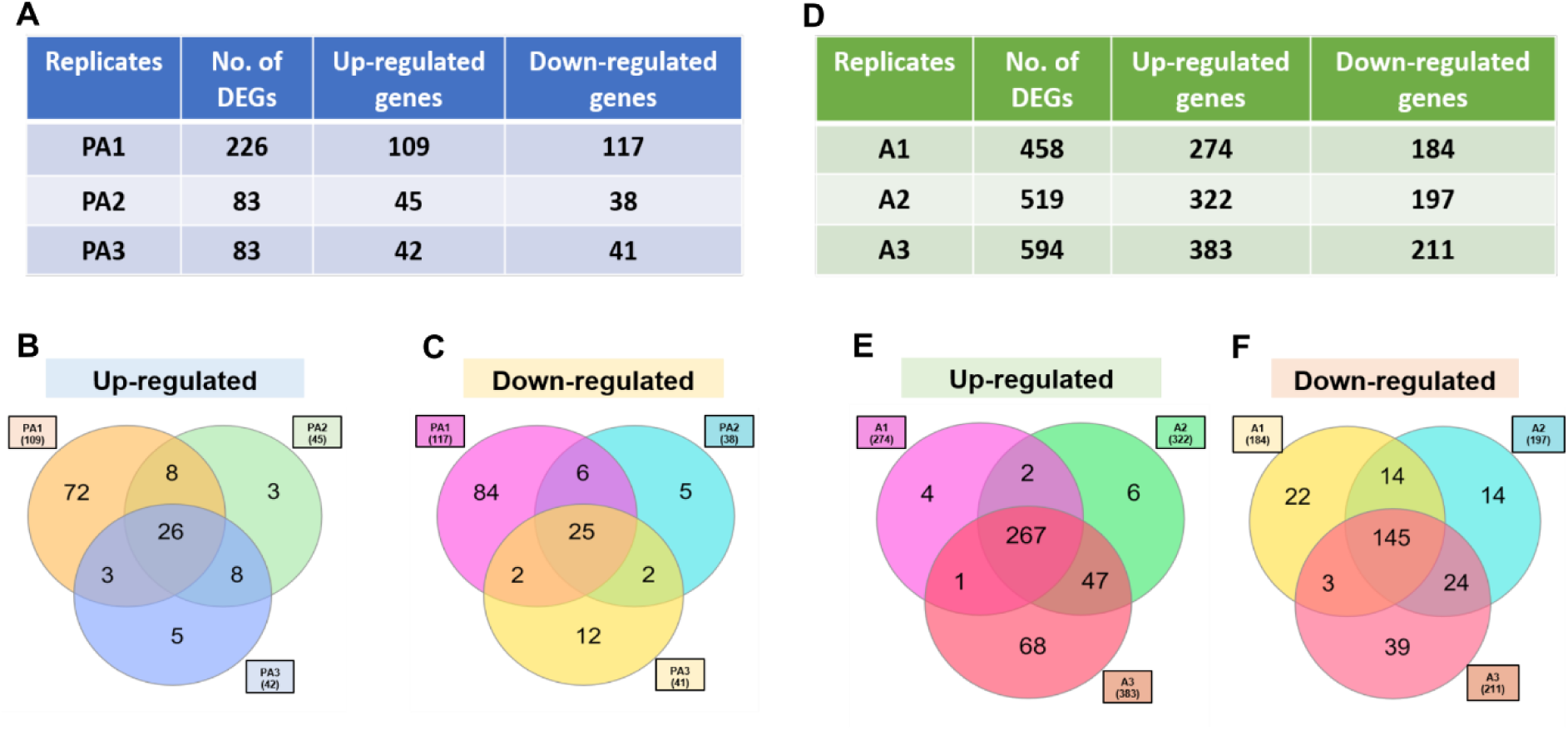
Global transcriptomics analysis of the adaptors of both P2428, and CBS10913^T^. P-line (left panel), and A-line (right panel) **A**, and **D,** tables depicting the total number of differentially expressed genes along with the up and down-regulated gene numbers. **(B, C), and (E,F)** showing Venn diagrams with the common Up and Down-regulated genes and adaptor exclusive genes in the adaptors of P2428, and CBS10913^T^.

### An alternative oxidase *AOX2* impacts AmB resistance in both A- and P-line adaptors

Our DEG analysis revealed significantly higher upregulation of the hitherto unknown gene *AOX2* encoding an alternative oxidase in AmB-adapted lines. We observed that AmB adaptors of A-line show 5-6 log_2_-fold overexpression of *AOX2* as compared to P-line adaptors, which showed 0.8-0.9 log_2_-fold increase in its expression (Fig. S5 A and B). The up-regulation of *AOX2* was associated with a significant decrease in ROS levels in both A - and P-line adaptors (Fig. 3A, B) and (Fig. S5 C and D). The involvement of *AOX2* in AmB resistance became more apparent when we examined the drug susceptibility and ROS levels in *AOX2*-deleted adaptor lines. For this, we deleted *AOX2* gene in one replicate of each (PA3 and A3) which exhibited highest MIC_50_ values along with their respective parental control strains (P2428 and CBS10913^T^). The knockout strains were designated as P2428/*aox2Δ*, and PA3/*aox2Δ* for the P-line, and CBS10913^T^/*aox2Δ* and A3/*aox2Δ* for the A-line adaptors. The growth kinetics of *aox2Δ* strains were comparable to native strain and deletion of *AOX2* did not lead to any growth alterations (Fig. 3A(iv). Notably, the *aox2Δ* in the adaptors (PA3/*aox2Δ* and; A3/*aox2Δ*) manifested several fold increased susceptibilities towards AmB when compared to their original progenitors (Fig. 3E, and F). For instance, PA3/*aox2Δ* showed MIC_50_ of 0.25-0.5 µgmL^-1^ in comparison with MIC_50_ of 16-32 µgmL^-1^ of parental controls. Moreover, A- and P-line adaptors showed significant corelation with ROS levels. But, in case of their respective *AOX2* deletants, unlike A3/*aox2Δ,* PA3/*aox2Δ* did not show good correlation with ROS levels. There was a collateral expected increase in ROS levels only in A3/*aox2Δ* cells while PA3/*aox2Δ* cells did not show any significant change (Fig. 3 C). The increased susceptibility towards AmB in *aox2Δ* strains was also supported by the growth kinetics of both the deletants when they were grown in the presence of higher concentrations of AmB (4 µgmL^-1^) (Fig. 3 E). Interestingly, the susceptibility towards AmB remained unchanged when *AOX2* was deleted in native progenitors CBS10913^T^ and P2428 (Fig. 3 F).

**FIG 3:**
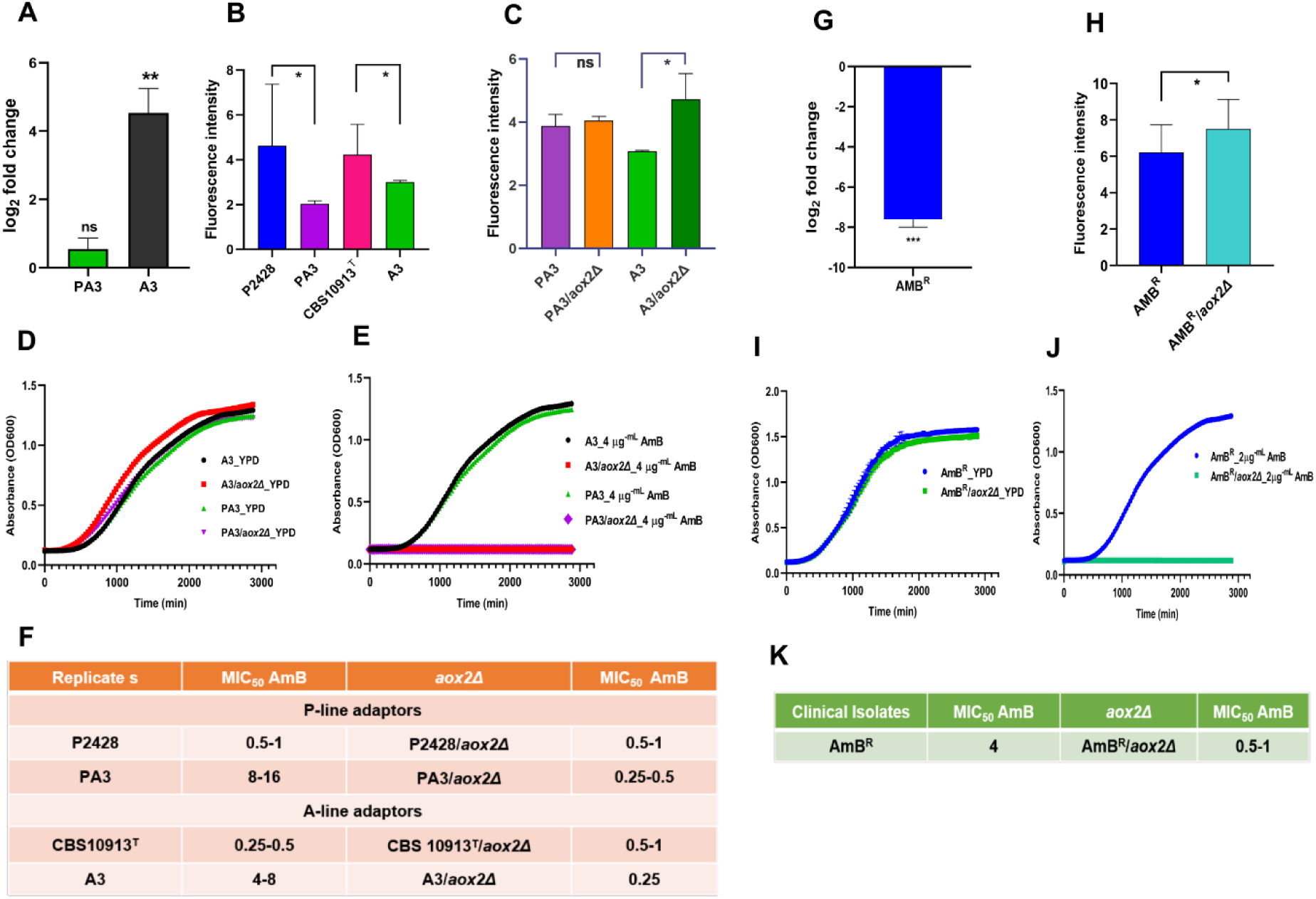
*AOX2* impacts AmB resistance. **A.** Left panel **A** The difference in expression levels of the *AOX2* gene in the adaptors, PA3 and A3, validated by SYBR green dye based Real-Time Quantitative PCR. The expression levels were normalised against the *C. auris* housekeeping gene *ACT1*. **B**, and **C,** The levels of ROS in the adaptors and in the respective *AOX2* deletants. The ROS levels were estimated by the fluorescent dye DCFH-DA. The cells were incubated with the dye for 30 minutes in dark followed by washing with PBS buffer and measuring the fluorescence with excitation and emission wavelengths 480 nm and 540 nm, respectively in a spectrofluorometer (Cary eclipse spectrofluorometer, Agilent USA). **D,** Growth pattern difference among the adaptors and their respective deletants in YEPD, and **E,** in the presence of 4 µgmL^-1^ AmB. All the experiments were performed in biological triplicates with technical duplicates. **F,** Table enlisting the MIC_50_ values of adaptors towards AmB before and after deletion of the *AOX2* gene. **G,** The difference in expression levels of the *AOX2* in the AMB^R^ clinical isolate validated by SYBR green dye based Real-Time Quantitative PCR. **H,** estimation of ROS in the AMB^R^ clinical isolate and in the respective *AOX2* deletant. The ROS levels were estimated by the fluorescent dye DCFH-DA as described above for Fig 3A. **I,** Growth kinetics of the AMB^R^ clinical isolate, the *AOX2* deletant in YEPD, and **J,** in the presence of 2 µgmL^-1^ AmB. All these experiments were performed in biological triplicates and technical duplicates. **K,** Table enlisting the MIC_50_ values of AMB^R^ clinical isolate towards AmB before and after deletion of the *AOX2* gene.

We also explored the role of *AOX2* in an AmB^R^ clinical isolate of clade I. Unlike AmB resistant adaptor lines, the expression level by qRT-PCR, revealed *AOX2* as among the most down-regulated genes (−6.7 log_2_-fold changes in NCCPF 470140 (AmB^R^) clinical isolate (Fig. 3 G). It should be pointed out that we also observed downregulation of *AOX2* in two more sets of AmB^R^ clinical isolates not included in this study (data not shown). Although, we adapted two drug susceptible isolates of clade II to AmB resistance, hence a strict comparison between AmB resistant clade I clinical isolate and clade II adaptors cannot be made. Nonetheless, the regulation of *AOX2* expression appears to be the opposite between clade II-AmB adaptors and AmB^R^ clinical isolate of clade I. The direct exposure of AmB as in adapter cells likely results in transcriptional activation of *AOX2,* which may not be the case with the clinical isolate. Nonetheless, the role of *AOX2* in impacting AmB resistance seems consistent with our observations from adaptor lines. For instance, the deletion of *AOX2* in AmB^R^ clinical isolate of clade I resulted in an increased susceptibility towards AmB in the AmB^R^/*aox2Δ* cells. (Fig. 3 K). This feature of AmB^R^ clinical isolate and its *AOX2* deletant was also evident in their growth patterns (Fig. 3 I, and 3 J). The ROS levels were further elevated in the AmB^R^/*aox2Δ* cells as compared to their progenitor (Fig. 3 H). Our data suggest that the alternative oxidase *AOX2* levels impact AmB susceptibility in the adaptors of clade II as well as in the resistant clinical isolate of clade I. However, its exact role still requires to be explored.

### AmB resistant adaptors show compromised cell wall integrity

The DEG analysis revealed that certain genes related to cell wall (CW) structural constituents (*PIR2*), and GPI-anchored CW-proteins (*PGA30*, and *PGA31*), were commonly upregulated in all three adaptors of P-line. On the other hand, an adhesin-like protein (*RBR3*) was commonly downregulated in these adaptors. In all the adaptors of A-line, genes associated with cellular glucan metabolic process (*XOG1*), cell surface glycosidase acting on CW 1,3-β-glucan 1,3-beta-glucanosyltransferase having role in fungal-CW (1,3)-beta-D-glucan biosynthetic process (*PGA4*), GPI-anchored adhesin-like proteins (*PGA7,* and *PGA48,* respectively) were downregulated. We validated the expression levels of common DEGs present in all the three adaptors of each progenitor by qRT-PCR as depicted in (Fig. 4A, and 4D). The impact of DEG related to CW synthesis was evident from the increased susceptibility of adaptor lines towards CW perturbing agents. We observed that both the sets of adaptors of A- and P-line display increased susceptibility towards Calcofluor White (CFW) and Congo Red (CR) (Fig. 4 C, and 4 F). Adaptor A3 of A-line was the only exception that showed resistance towards both the CW perturbing agents. Additionally, all three adaptors of P-line exhibited resistance towards FK506, which is a calcineurin inhibitor (Fig. 4 B). In contrast, replicates of A-line remained susceptible towards FK506, where A3 was the only exception displaying resistance towards the calcineurin inhibitor (Fig. 4 E). For the chitin synthesis in fungal CW, three pathways are supposed to be cumulatively participating including HOG MAP-kinase pathway, PKC-signalling cascades, and Ca^2+^/Calcineurin-pathway (22). Calcineurin also plays an important role in attenuating the excess synthesis of chitin (23). In the present study, all the adaptors of P-lines display resistance towards FK506, implying activation of calcineurin. As a result, the cells are sensitive towards CFW. Whereas in A-lines, except A3 replicate, the CFW susceptible adaptors are also sensitive towards FK506 (Fig. 4). This highlights the intra-clade differential impact on CW integrity (CWI), and its role therein towards the development of drug resistance mechanisms. Additionally, the *MSN4* gene can also act as a chitin synthesis inhibitor (23). This gene was downregulated in all the adaptors of A-line. So, specifically in case of A3, the *MSN4* gene might have an effect as this adaptor is resistant to CW-perturbing agents. (Fig. 4).

**FIG 4:**
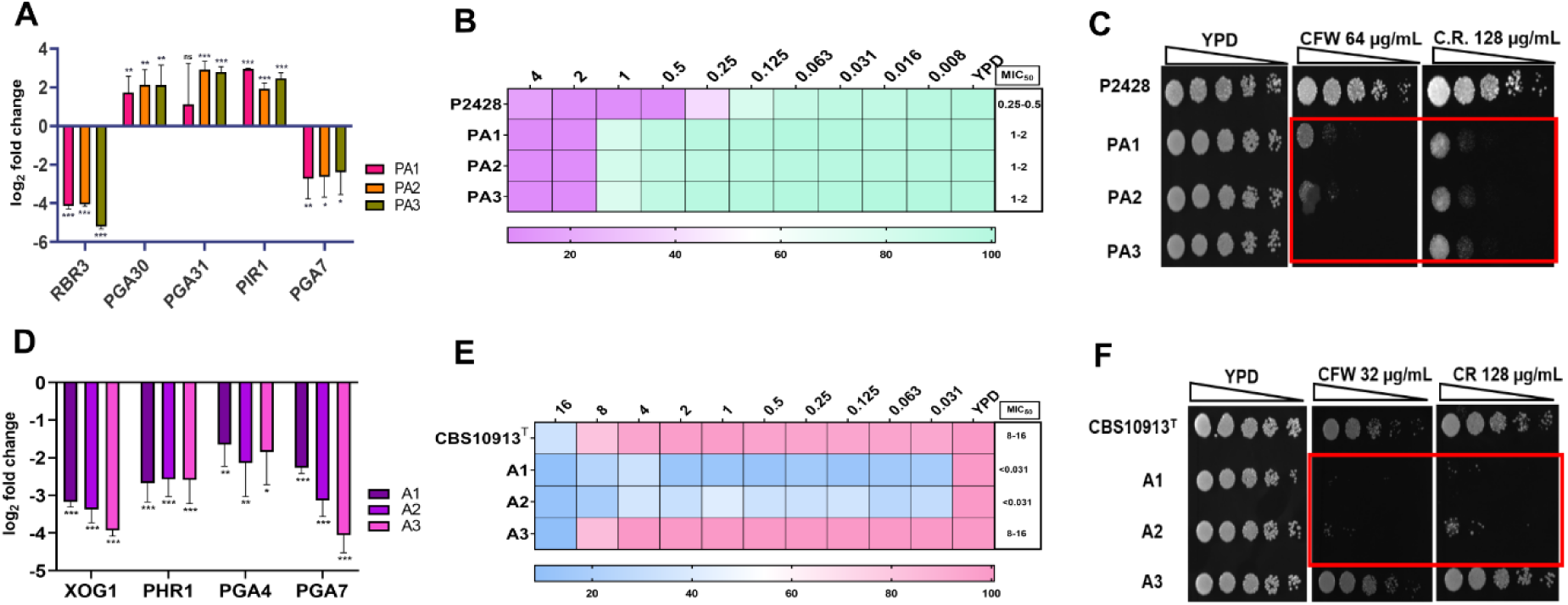
Characteristics of the cell wall in the adaptors of A and P-lines. **A**, and **D,** Adaptor lines of P-line, and A-line displayed increased susceptibility to CW perturbing agents. Spot assays depicting susceptibility towards CFW and CR for P, and A-adaptor lines. The OD_600_ of overnight grown cells was set to 0.1 and serially diluted up to five dilutions, which were then spotted onto YEPD plates (3 µL of spot volume) with or without the drug. Growth differences were recorded after 48 hrs of incubation at 30°C by BioRad XR+ Gel documentation system. **B,** and **E** depict the grid showing susceptibility towards FK506 tested by broth microdilution assay. Cells were evaluated for growth in the presence of FK506 post 48 hrs incubation at 30 ⁰C in BioRad iMark microplate reader. **C,** and **F**. The expression of CW-related genes was validated by Real-Time Quantitative PCR normalised against the housekeeping gene *ACT1*.

### A-line adaptors can drive AmB resistance independent of *ERG* genes

The global transcriptomic analysis revealed interesting insights of the evolvability of two different *C. auris* isolates of same clade in acquiring high resistance to AmB. The differential expression or mutations in *ERG2*, *ERG3*, *ERG6, ERG11* and *ERG25*, which leads to defect in ergosterol biosynthesis are also shown to impact the AmB susceptibility (10); (9, 11, 24, 25). However, our present stepwise evolution study points to the potential intra-clade-heterogeneity in evolving to AmB resistance. For instance, AmB adapted P-lines which displayed 16-32-fold increase in MIC_50_ values, the ergosterol biosynthesis pathway genes such as *ERG1, ERG2, ERG4, ERG5, ERG6*, *ERG10, ERG13, ERG24,* and *ERG25* were among the upregulated genes in most of the replicates (Fig. 5A). That is also reflected in increased ergosterol levels in adapted cells (Fig. 5 B). However, no such correlation between the expression of *ERG* genes and ergosterol levels was observed in A-line adaptors. The A-line adaptors, which also showed a 2-4 folds increase in MIC_50_ value towards AmB, did neither displayed any differential expression of the *ERG* genes nor any increase in ergosterol levels. The ergosterol levels in A-line adaptors were rather reduced in all the adapted cells (Fig. 5 D).

**FIG 5:**
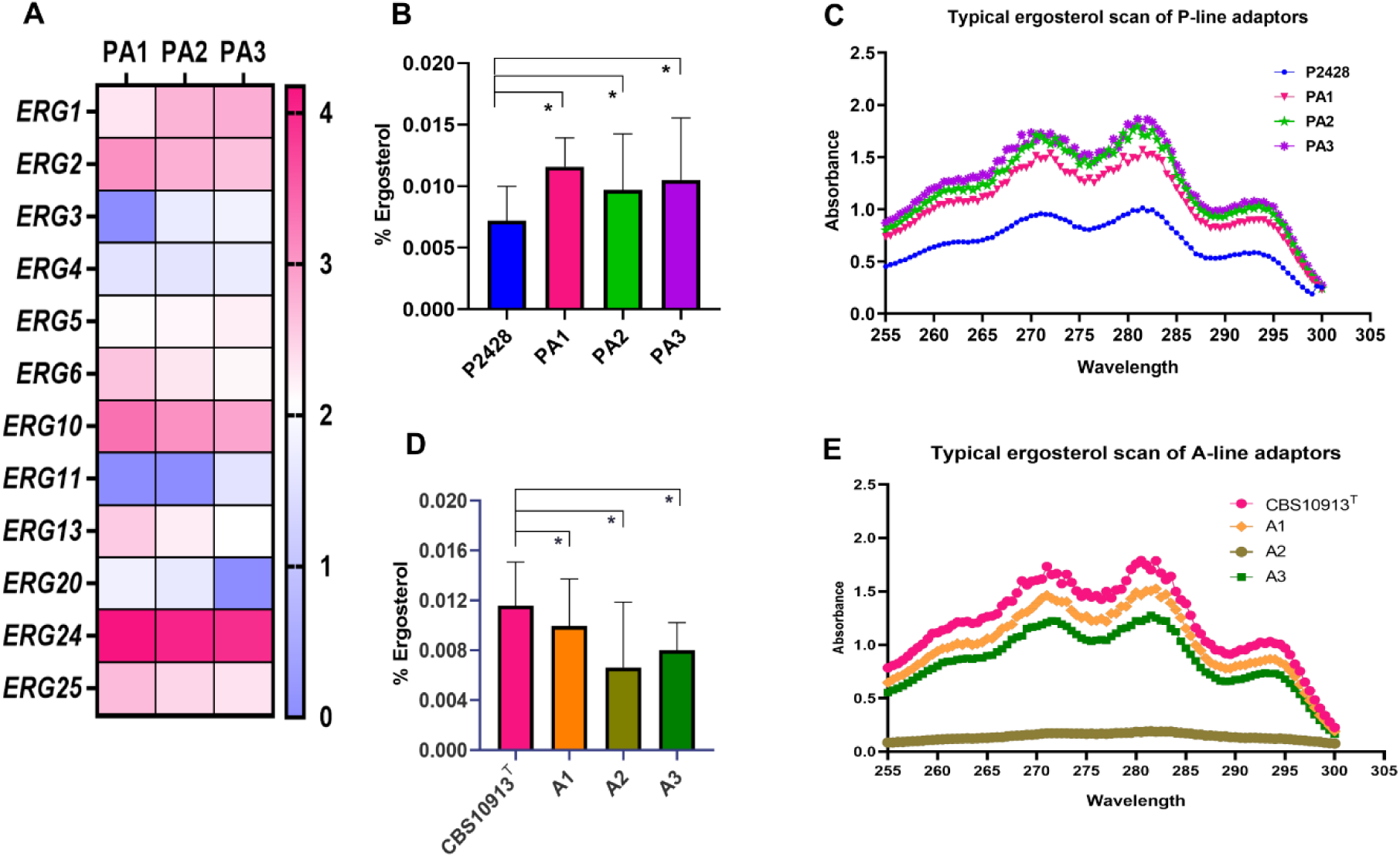
*ERG* genes expression and the ergosterol content in the adaptors. **A**. Heatmap depicting the differential expression levels of the ergosterol biosynthesis pathway genes observed in the RNA sequencing results of P-line adaptors. **B,** Ergosterol percentage in P-line adaptors. The ergosterol percentage was calculated by calculating the ratios of ergosterol and 24(28)-dehydroergosterol DHE by the absorbance values obtained at 281.5 nm and 230 nm for ergosterol and DHE, respectively. **C,** A typical ergosterol scan of the P-line adaptors. **D,** Ergosterol percentage in A-line adaptors. **E,** A typical ergosterol scan of the adaptors of A-line. All these experiments were performed in biological triplicates, the means and standard deviations were then calculated and plotted.

### Glucosylceramides were absent in P line adaptors

Among DEGs of the P-line adaptors, the gene *HSX11*, which encodes glucosylceramide (GlcCer) synthase, was found to be highly downregulated (−4.33-log fold change) and was commonly affected in all three adaptors. *HSX11* catalyses the transfer of UDP-linked glucose to the sphingoid backbone of ceramide precursors. To confirm the impact of this downregulation at the metabolic level, we conducted mass-spectrometry-based sphingolipid analysis. The results showed a significant reduction of GlcCer levels in P-line adapted strains (<1%) compared to the control (∼60%). As expected, there was a parallel accumulation of precursor of GlcCer, which is α-hydroxyceramides, in all the adapted cells (Fig. 6 B). Significantly, such downregulation of *HSX11* gene and the corresponding decrease in GlcCer levels was not evident in A-line adaptors. While, we detected several molecular species with different sphingoid backbones, such as d18:1, d18:2 and d19:2 and different fatty acid chains, the major GlcCer contributing species was GlcCer (d19:2/18:0(2OH[R])) and its precursor Cer (d19:2/18:0(2(OH[R])) was accumulated in the adapted P-lines (supplementary file S9). Based on our earlier analysis, glucosyl derivatives are the main complex sphingolipids (SLs) found in *C. auris*, indicating that the GlcCer branch of the SL pathway is active in this species (26). However, the specific role of GlcCer levels in drug resistance is still not fully understood. We observed a significant reduction in GlcCer levels in the P-line adaptors, but further investigation is needed to determine its role in AmB resistance. In our WGS analysis, a common SNP was detected in all the adaptors of the P-line, *B9J08_000315 ^K122R^*, which is an ortholog of *S. cerevisiae CSG2* which codes for ER-calcium channel and is required for mannosylation of IPCs. Inositol phosphorylceramides (IPCs) are a class of complex anionic sphingolipids found in fungi plants and some protozoans but absent in mammals. These are characterized by presence of Inositol group linked to Ceramide backbone at C-1 position. In fungi are known to interact with sterols in the form of lipid rafts and are known to mediate multiple roles including virulence (27). Our lipid analysis of P-line adaptors could only quantify IPCs and observed their reduction in the adapted lines implying an increase in mannosylated IPCs (MIPC/M(IP)_2_C (Fig. 6 B). Notably, spot assays of the P-line adaptors showed increased susceptibility in YPD media supplemented with 1M CaCl_2_ as compared to their parent, P2428 cells (Fig. 6 E).The increased susceptibility towards CaCl_2_ and decrease in IPCs could reflect the impact of missense mutation in the ortholog of *CSG2* gene. Interestingly, we also found that several other genes related to iron metabolism (*B9J08_000008*), phosphatidylinositol transfer (*B9J08_000007*), carnitine acetyl transferase (*B9J08_000010*), and an uncharacterized gene (B9J08_00006) were co-downregulated in the same scaffold PEKT02000001 where *HSX11* is present (Fig. 6 A). These genes are likely to be regulated by a common transcription factor, which may be affected during the development of resistance to AmB. Among the common genes exhibiting SNPs in all three P-line adaptors, *B9J08_005579 (HAP41)* was a regulator which was affected commonly. However, it is not yet confirmed whether *HSX11* and other genes in the same scaffold are part of *HAP41* regulon. Interestingly, in A-line adaptors GlcCer levels remain unchanged highlighting another instance of intra-clade-heterogeneity.

**FIG 6:**
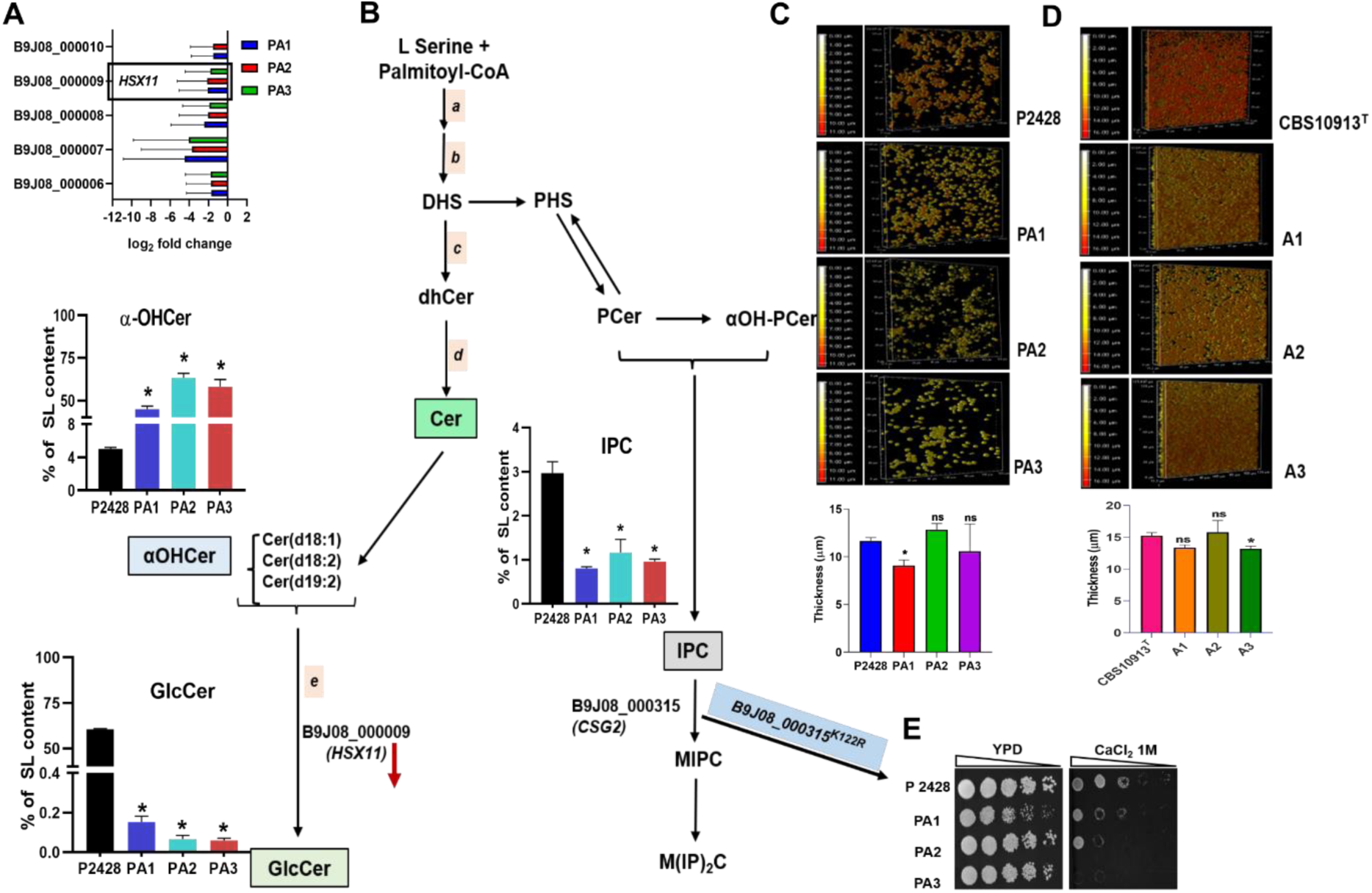
Sphingolipidomics of in P2428 adaptor lines. **A.** The expression level of *HSX11* (glycosyl ceramide synthase) in P-line adaptors. The expressions of genes in the same scaffold along with *HSX11* is also depicted in the panel. These genes were co-downregulated along with the *HSX11* gene present in the scaffold, PEKT02000001. **B**. Schematic representation of part of the Sphingolipid biosynthesis pathway depicting the steps leading to the synthesis of GlcCer and the levels of the intermediates shown as bar charts along with the pathway. The enzymes catalysing various steps of the pathway are; a) Serine palmitoyl transferase, b) 3-Keto dihydro Sphingosine reductase, c) Ceramide synthases, d) Δ4-desaturase, and e) GlcCer synthase (*HSX11*). **C**, and **D** Biofilm formation study of the AmB adaptors of **C,** P-line, and **D,** A-line. **E,** Spot assay grid of P-line adaptors with CaCl_2_.

### AmB adapted cells show fragile biofilms

Adhesion is the first important step in biofilm development. Adhesins are glycosyl-phosphatidylinositol-cell wall proteins (GPI-CWPs) that are composed of a GPI anchor, a serine/threonine domain, and a carbohydrate or peptide binding domain (28). Most of the CW assembly related genes found in the present study are GPI anchors which are downregulated, for instance *PGA7* and *RBT5* are commonly down regulated in both A and P-line adaptors. Being a crucial step, the downregulation of these genes indicates less adhesion ability of the biofilms which supports the observation that all adaptors of A and P-lines in general form very thin and fragile biofilms as compared to their respective parental strains (Fig. 6 C, and D). The β-1,3-glucan of the biofilm matrix have has the ability to particularly bind with the AmB and thereby diminishing its effect on fungal cells encased in the biofilm matrix (29). In a study conducted by Kean et al (2018) (30) in *C. albicans*, seven genes related to biofilm production and antifungal resistance were found to be upregulated across isolates of *C. auris, C. haemulonii, C. duobushaemulonii,* and *C. pseudohaemulonii*. In contrast to their observations, in the present study, we found only *PGA7* as a commonly downregulated gene in both the sets of adaptors. And the biofilms of all these adaptors are fragile and thin as compared to their respective controls. According to a previous study conducted by Kean *et al*, the elevated expressions of adhesins-related GPI-anchored CW genes are required for biofilm formation. In the present microevolution study, these genes’ downregulation might coincide with the observed fragile feature and thin biofilms. The reduced expression of these genes in the adaptors of both the lines leads to compromised CWI which is also reflected in enhanced susceptibility towards CW perturbing agents. A study conducted by (31) on filamentous fungus, *Scedosporium aurantiacum* explained the crucial role of GlcCers in growth, germination and pathogenicity. In that study, antibodies used against GlcCers reduced the biofilm adherence, biomass, and viability of the biofilms. In our analysis where the GlcCer are totally vanished from P-line adaptors, the biofilms are also too thin and fragile (Fig.6 C, and D). However, in A-line adaptors where GlcCer does not seems be crucial, the fragile biofilm could be due to yet other unknown contributors.

### Evolved AmB resistant A and P-lines show no change in ploidy

Karyotype analysis of the evolved A- and P-lines was conducted to assess chromosomal changes resulting from experimental evolution with AmB. No significant karyotype variations were observed in the adaptors compared to the parental controls (Fig. S6). This is in contrast to our previous findings on fluconazole-resistant isolates (32). Additionally, copy number variation analysis of DNA sequencing raw reads did not reveal any alterations in comparison to the parental genome.

### The Whole genome sequencing analysis unveils an intra-clade heterogenous landscape of SNPs

To understand the changes that could occur at the genomic DNA sequence level, whole genomes of the P- and A-line of adaptors were subjected to WGS along with their control strains. To identify non-synonymous mutations, a comparison was made between the adaptors and their respective control strains. In case of the P-line analysis, the adaptors were compared with the control strain P2428. Similarly, for A-line analysis, the adaptors were compared with the control strain CBS10913^T^. This approach allowed for the identification of specific mutations present in the adaptors of each line. Our analysis revealed that the number of genes showing SNPs were variable between the two lines (Fig. 7A, and 7C). For instance, in P-line adaptors, total number of genes exhibiting missense mutations observed in PA1, PA2, and PA3 were 257, 118, and 127, respectively. Among these genes, 232 were exclusively present in PA1, 61 in PA2, and 72 were exclusive for PA3 (Fig. 7 A). Together, there were 9 common genes with SNPs among all the three P-line adaptors (Fig. 7 B). Unlike, in A-line adaptors, the number of genes having SNPs was lower. For instance, there were only 19, 20, and 23 genes having missense mutations in A1, A2, and A3 adaptors, respectively. Among them, only 4 genes were present exclusively in A1, 3 in A2, and 9 in A3 (Fig. 7 C). Despite low number of SNPs in A-line adaptors, they show 10 common genes with SNPs among all the three adaptors (Fig. 7 D). Among all the adaptors of both the lines, only one uncharacterised gene, *B9J08_005550* (*orf19.5281*) commonly exhibited missense mutations. In this gene, there were two SNPs in P-line adaptors and one different SNP in A-line adaptors (Fig. 7). This ORF remains uncharacterised in *C. auris* but its orthologue in *C. albicans* has a role related to nuclear envelope and endoplasmic reticulum (33). One more study also hypothesized that Scp160p participates in cytoplasmic mRNA metabolism, which may encompass translation, though the exact biological role of Scp160p remains undefined (34). In the adaptors of both the lines, the SNPs present are from different categories like regulation of biological process, transport, RNA metabolic process, organelle organization, response to stress, response to chemical, DNA metabolic process, lipid metabolic process, cellular homeostasis, cell wall organization, cell development etc (Fig. S7).

**FIG 7:**
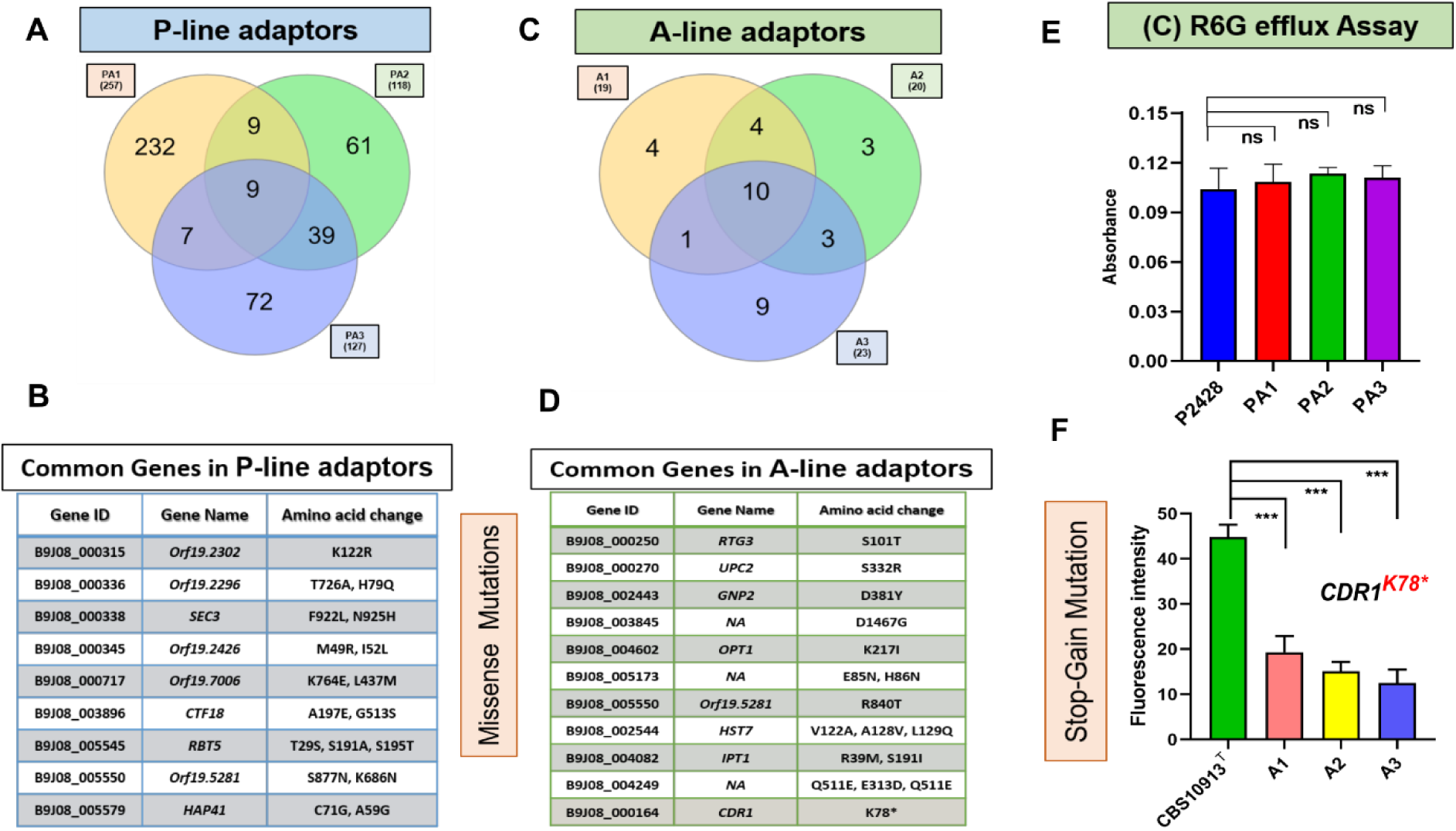
Landscape of SNPs in A- and P-lines. Venn diagrams **A**, and **C,** explaining the total number of genes exclusive for the individual adaptors and common genes among the adaptors exhibiting missense mutations. and lists of common SNPs present in the adaptors of **B,** P-line, and **D,** A-line. R6G efflux assay in **E,** P-line. and **F,** A-line. Exponential phase adaptor cells were incubated with R6G for 3 hours. Post incubation the cells were washed with PBS buffer and divided into two sets. One set was provided with 2% glucose for further 45 minutes, whereas another set was kept devoid of glucose for energy depletion. After the incubation with/without glucose, the cells were washed with PBS and resuspended in 1 mL PBS buffer. The fluorescence was measured at excitation and emission wavelengths of 440 nm and 570 nm wavelengths respectively by Cary eclipse spectrofluorometer, Agilent USA. The experiment was carried out in biological triplicates with technical duplicates.

**FIG. 8:**
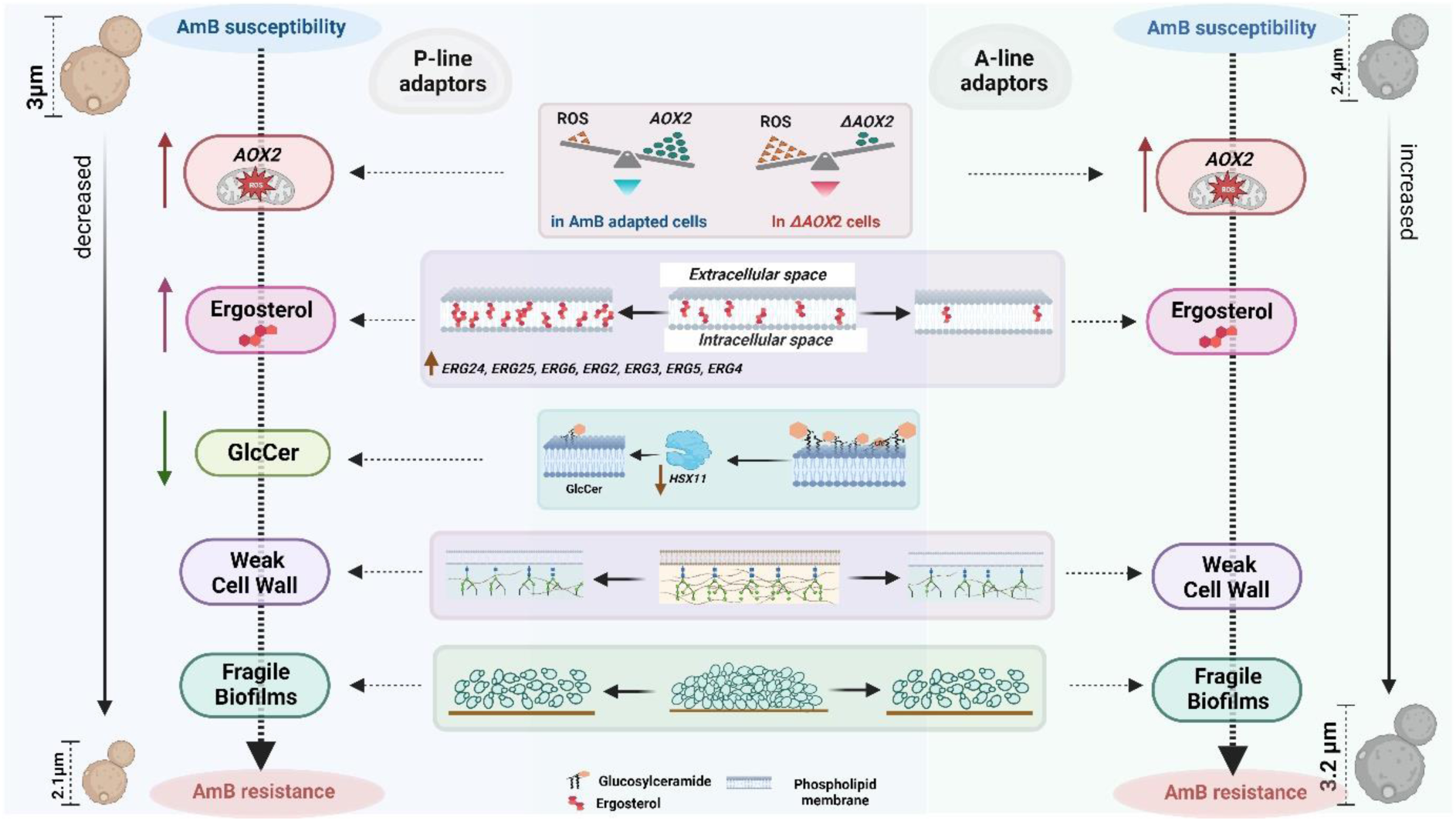
Schematic depiction of various factors imparting AmB resistance in the P2428 and CBS10913^T^ lines of adaptors.

Among 9 common genes with SNPs in P-line, 5 were in uncharacterized ORFs and the rest were in *B9J08_005545 (RBT5^T29S, S191A, S195T^), B9J08_005579 (HAP41^C71G, A59G^), B9J08_003896 (CTF18^A197E, G513S^), B9J08_000338 (SEC3^F922L, N925H^)* genes (Fig.7 B).When categorized by their association with CW functions, *RBT5* displayed a consistent pattern of downregulation in all adaptors of both lines. Furthermore, *RBT5* was found to harbor three common SNPs at *RBT5^T29S, S191A, S195T^*.

The *erg6* mutant of *C. neoformans* and *C. glabrata* both show sensitivity to CR and SDS. This sensitivity also suggests possible disturbances in protein trafficking or mis-location of CW-related enzymes (10, 35). The *erg6* mutants are sensitive to azoles and allylamines, but resistant to polyenes (9). In the P-line adaptors, most *ERG* genes do not have missense mutations, but two adaptors (PA2 and PA3) have a new mutation in *ERG6*^L58V^ and a novel SNP in *UPC2*^I345L^. Interestingly, in adaptor PA3, the expression of *ERG11* is also upregulated, which may be related to the novel SNP in *UPC2*. These observations suggest a potential role of *ERG6* and *UPC2* in influencing resistance to AmB in P-line adaptors. This does not, however, seem to be the case in A-lines, which exhibit ergosterol independent resistance to AmB. The occurrence of a new SNP in *UPC2^S332R^* was detected in all A-line adaptors, but it does not appear to have any functional consequence. There were 10 common genes in A-line adaptors which harboured SNPs (Fig. 7 D). These include, *B9J08_000270 (UPC2^S332R^), B9J08_000250 (RTG3^S101T^), B9J08_002443* (*GNP2^D381Y^*), *B9J08_003845, B9J08_004602 (OPT1^K217I^), B9J08_005173, B9J08_005550 (orf19.5281^R840T^), B9J08_002544 (HST7^V122A, A128V, L129Q, A130Q^), B9J08_004082 (IPT^R39M, S191I^), B9J08_004249*.

It is interesting to note that among the transporter genes with SNPs, a notable finding is the presence of a nonsense mutation in the major drug transporter *CDR1* in all the A-line adaptors. This mutation (*CDR1^K78^*\*) is expected to result in a non-functional truncated Cdr1 protein. Although we did not directly measure the levels of Cdr1p in our adaptor cells, we assessed the functional status of this efflux pump. The efflux of Rhodamine 6G (R6G), a substrate of Cdr1p, was found to be reduced in all the A-line adaptors compared to their parental strain, CBS10913^T^ (as shown in Fig. 7 F). In contrast, there was no change in the efflux of R6G and no SNP detected in P-line adaptors (Fig. 7 E). Given the presence of a significant number of genes with SNPs among the adapted lines within the same clade, it is crucial to conduct thorough validation to determine their functional significance. This validation process will help us understand the specific roles and effects of these genetic variations (as shown in Fig. S7) and gain a better understanding of their impact on the organism.

## Discussion

Our study aimed to understand the mechanism underlying the widespread amphotericin B (AmB) resistance in *C. auris*. We selected two drug-susceptible *C. auris* cells from different geographical locations: CBS10913^T^, isolated from a patient’s ear canal in Japan, and P2428, recovered from an Indian diabetic patient’s pus. These isolates underwent directed evolution by continuous exposure to sub-lethal concentrations of AmB for 100 generations. This approach allowed us to investigate the specific alterations that occurred during the evolution of AmB resistance in *C. auris* isolates from different backgrounds yet belonging to same clade. Our findings not only shed light on the intricate processes that contribute to AmB resistance but also signify intra-clade heterogeneity which exists in *C. auris*. Apart from its well-established interaction with fungal sterol, the mechanisms leading to the fungicidal effects of AmB are highly complex. Extensive evidence suggests that AmB’s action extends beyond merely binding and depleting membrane ergosterol. In fact, there is a substantial body of research indicating the involvement of oxidative stress in AmB’s fungicidal effects. AmB can undergo autoxidation, thus causing oxidative stress. This interplay between AmB and oxidative stress plays a crucial role in its effectiveness against fungi (36). Several reports suggest that hypoxia can provide protection to protoplasts of *C. albicans* cells against AmB. It has been observed that the addition of catalase and superoxide dismutase (SOD) can effectively prevent the AmB-induced lysis of protoplasts. Similarly, *Aspergillus terreus*, which is intrinsically resistant to AmB, has been found to exhibit a significant increase in catalase levels without any notable changes in ergosterol levels (37). This suggests the presence of a catalase-based resistance mechanism that counteracts the oxidative stress induced by AmB, ultimately preventing cell damage. (38, 39).

While previous studies have indicated the involvement of *AOX2* in mitochondrial alternate respiration, mycelial development, and virulence in *Candida* cells, there is currently no direct evidence suggesting its involvement in AmB resistance. However, our present study has made a direct observation regarding the influence of *AOX2* on AmB susceptibility through experimental evolution. The fact that *AOX2* was among the most upregulated gene in our AmB resistant adaptor lines prompted us to examine the impact of it by deleting *AOX2* in AmB-resistant adapted lines of *C. auris* from both A and P-line adaptors. The resulting *aox2Δ* cells of both the lines exhibited enhanced susceptibility to AmB. This finding highlights the significant impact of *AOX2* in modulating the susceptibility of *C. auris* to AmB. Interestingly, there was a significant drop in ROS levels in both the adaptor lines, coinciding with the upregulated expression of *AOX2* gene in these lines. This suggests that the enhanced levels of *AOX2* may contribute to better management of ROS in AmB-resistant lines. However, it is important to note that the direct correlation between ROS and AmB resistance was not evident in all cases, indicating that alternate mechanisms may be at play. Additionally, *AOX2* appears to be a specific feature of A-line adaptors, as it is highly upregulated as compared to that in the P-line adaptors. This suggests that the evolution of AmB resistance may still be heterogeneously manifested within the isolates of same clade. Further investigation is needed to establish a conclusive relationship between altered ROS levels and AmB susceptibility in *C. auris*.

Our data has established a compelling correlation between compromised CWI and AmB resistance in adaptors of both lines. This significant observation provides further evidence supporting the notion that oxidative stress has a direct impact on CWI which is evident from increased susceptibility towards CW perturbing agents along with differential expression of CW related genes in *C. auris*. In our current experimental evolution study, we have observed that the adaptor lines exhibit downregulation in the glycolysis pathway, indicating a metabolic shift. Despite this compromised effect on the respiratory pathway, the adaptors remain resistant to AmB. This suggests that the cell employs alternative respiration as a survival mechanism during AmB stress, leading to the elevated expression of *AOX2*. This significant observation provides further evidence supporting the idea that oxidative stress directly affects CWI in adaptor lines of *C. auris* belonging to clade II.

Our findings suggest a contradiction to the conventional observation regarding the impact of ergosterol levels on AmB resistance in clade II adaptors. Surprisingly, P-line adaptors exhibited upregulation of essential *ERG* genes and an increase in ergosterol levels, while no such changes were observed in the AmB-resistant lines of A-series. This observation challenges the long-standing understanding of the relationship between ergosterol and AmB resistance in *C. auris* and other fungi, indicating that alternative mechanisms may be at play in conferring resistance within clade II. We also observed isolate-specific perturbation in sphingolipid metabolism, with P-line adaptors showing down-regulation of *HSX11*, a gene encoding glycosyl ceramide synthase, resulting in a significant loss in GlcCer and an accumulation of its precursor OHCer. However, A-line adaptors did not show any difference in *HSX11* expression or sphingolipid composition (Supplementary file S9). These findings suggest that the P-line adaptors rely more on lipid metabolism as compared to the A-line. These novel insights provide exciting avenues for further investigation into the complex interplay between ergosterol, sphingolipids, and AmB resistance in different clades of *C. auris*.

Our microevolution study resulted in the identification of a new set of SNPs that have not been previously reported. In a study by (16), they observed mutations in 8 genes (*ERG3^W182*^, ERG11^E429*^, MEC3^A272V^, FLO8^Q384*^, FKS1^FL635L^, CIS2^A27T^, PEA2^D367V^*, and a duplication event Chr1^Pdup^ during the microevolution of *C. auris* towards AmB, FLC, and Caspofungin resistance. However, in our microevolution study, we found a completely different set of mutations in the AmB adaptor lines that did not include any of the genes mentioned above. Comparably, we found SNPs in *RBT5^T29S, S191A, S195T^*, *PGA7^G185D^, MKC1 ^I368T^, CEK1^S122N^, HST7^N478K^, ZCF14^V781I^, TIM21^L76F^, SIT1^R74K^, LIP1^Q25E^, INO1^M16^* in the P-line adaptors. And in the A-line adaptors, among the genes exhibiting SNPs were, *GNP2^D381Y^, CWH41^F400D^, IPT1^R39M^, OPT1^K217I^, ALS4^L115V^,* and *INN1^T194K^*. While, the relevance of these missense mutations in AmB adapted lines remains to be studied, it signifies the complicated nature of AmB resistance in *C. auris*.

## MATERIALS AND METHODS

### Strains and media

We obtained C. *auris* clade II isolate (CBS10913^T^) from the Central bureau voor Schimmel Cultures (CBS), Fungal Biodiversity Centre of the Royal Netherlands Academy of Arts and Sciences (KNAW), Utrecht. Another Clade II susceptible isolate (P2428) was procured from National Culture Collection of Pathogenic Fungi (NCCPF), Indian Council of Medical Research (ICMR), New Delhi, sponsored National facility at the Mycology Division, Department of Medical Microbiology, Postgraduate Institute of Medical Education and Research (PIGMER), Chandigarh, India. The ancestor strains and the evolved strains of every transfer were stored in −80 °C in 50% glycerol and, when used, then grown in YEPD (1% yeast extract, 2% peptone, 2% dextrose) at 30°C. All the strains used in this study are listed in Table S1.

### Clade-typing

The clade-specific primers reported previously were used to identify the clade-status of the isolates (20). The whole genome data was also compared with the available GenBank assemblies of strains belonging to different clades.

### *In vitro* evolution of *C. auris* clade II susceptible isolates P2428 and CBS10913^T^

For *in vitro* evolution, *C. auris* strains CBS10913^T^ and P2428 (MIC_50_-AmB-0.5 µgmL^-1^, and 1 µgmL^-1^) was used. The protocol of *in vitro* evolution described earlier by (40) and our group (32), was followed with some modifications. The strains were revived from a frozen stock on a YEPD plate and incubated for 24 hours at 30°C. Subsequently, a single colony was patched on a new YEPD plate and incubated for another 24 hours at 30°C. A single colony from this plate was cultured in a fresh 10 ml YEPD broth and incubated for 72 hrs at 30°C. From the stationary culture, 10 µL of 0.1 O.D._600_ cells were transferred into three independent tubes, each containing 9990 µL fresh YEPD with or without AmB (1 µgmL^-1^ and 0.5 µgmL^-1^ of AmB (for P2428 and CBS10913^T^, respectively) in triplicates in two different sets, one set as a control of replicates which was not exposed to the drug and were labelled as C1 and C2. The other set of replicates was continuously exposed to the AmB and were designated as A1, A2, and A3). Initially all the replicates and controls were incubated for 72 hours at 30°C. The culture from each replicate and control was transferred into fresh YEPD broth (with or without the drug, 10 µL from the previous culture in 9990 µL fresh YEPD broth resulting in 1:1000 dilution) and incubated for another 72 hours at 30°C. After 72 hours (10 generations), adapted cells with positive control were transferred into fresh media again in a 1:1000 dilution. One such transfer of 1:1000 dilution at 30°C for 72 hours corresponds to 10 generations (log_2_(1000) = 9.97, 1 transfer is equivalent to approximately 10 generations) (40). For 100 generations, 10 such transfers of the cells were performed in presence and absence of AmB. After every 10 generations, an aliquot of cells was drawn and stored at −80°C in 50% glycerol for further analysis.

### Gene deletion

*AOX2* was disrupted by homologous recombination with a cassette-containing *NAT1* gene, coding for nourseothricin acetyltransferase (imparts resistance to nourseothricin) flanked by 5′- and 3′-UTR regions of the gene. The 5′- and 3′-UTR (nearly 500 bp) of the genes were amplified from wild-type genomic DNA. Both the 5′ and 3′-UTR were fused to one-half each of the *NAT1* gene amplified from a plasmid.

The two *NAT1*-amplified fragments share an ∼300–350-bp complimentary region. The fused PCR products were co-transformed into the wild-type strain, and the transformants were plated on YPD. After incubation at 30°C for 12–16 h to allow homologous recombination within the *NAT1* fragments and with the genomic loci, cells were replica-plated onto a YPD plate supplemented with 100 µgmL^-1^ nourseothricin and further incubated for 24 h. Nourseothricin-resistant colonies were purified and checked for gene disruption via homologous recombination by PCR.

### Growth assays

The growth kinetics assays were performed by a micro-cultivation method in 96-well plates using Liquid Handling System (multimode microplate reader, Tecan, USA) in YEPD broth at 30°C. Briefly, overnight grown yeast cultures were diluted to 1.0 O.D._600_ and 20 µL of each culture was mixed with 180 µL YEPD broth with the selected antifungal AmB concentrations in a 96-well plate. O.D._600_ was measured at every 30 minutes for a period of 48 hours.

### Determination of Minimal inhibitory concentrations (MICs) and spot assays

MICs assay followed was of CLSI M27-A3 with some minor modifications. *C. auris* cells were grown overnight at 30°C in YEPD media and diluted in 0.9% saline solution to obtain an O.D._600_ nm of 0.1. The cells were then diluted 100-fold in YEPD medium. The diluted cell suspension was added to the wells of round-bottomed 96-well microtiter plates containing equal volumes of media and serially diluted concentrations of the drug. The plates were incubated at 30°C for 48 hours. The MIC test endpoint was evaluated by measuring the optical density at 600 nm in a microplate reader (Bio-Rad iMark) and was defined as the lowest drug concentration that gave 50% inhibition of growth (MIC_50_) as compared with the growth of the drug-free control.

### RNA Isolation

A saturated, overnight culture was used to inoculate 10 mL of fresh YEPD at an O.D._600_ of 0.1, which was grown for 4-5 hours at 30°C to obtain a log phase culture. The cells were then collected by centrifugation and washed with DEPC-treated water. Total RNA was isolated using RNeasy Mini Kit (Qiagen, Hilden, Germany, Cat No 74104), following the manufacturer’s specifications. Total RNA in the samples was quantified using Nanodrop 2000 spectrophotometer (Thermo Scientific, USA).

### RNA Sequencing and analysis

#### Sequence Data QC

The sequence data was generated using Illumina NovaSeq 6000. FastQC (https://www.bioinformatics.babraham.ac.uk/projects/fastqc/) was used to process the raw reads for quality assessment and pre-processing, which includes removing the adapter sequences and low-quality bases (<q30) using TrimGalore3 (https://www.bioinformatics.babraham.ac.uk/projects/trim_galore/). The pre-processed high-quality data were aligned to the reference genome (Candida Genome Database, *Candida auris* strain B8441) using Bowtie24 (41) with the default parameters to identify the alignment percentage. Reads were classified into aligned reads (which align to the reference genome) and unaligned reads. HTSeq5 (42) was used to estimate and calculate gene abundance. Absolute read counts for genes were identified and used in differential expression calculations. DESeq6 was used to identify the differentially expressed genes. Genes were categorized into up, down, and neutrally regulated based on the log2 fold change cut-off of 1. DESeq (43) normalized expression values were used to calculate fold change for a given gene. The regulation for each gene was assigned based on log2 fold change. The genes which show log2 fold change less than −1.5 are represented as down-regulated, the values greater than +1.5 are represented as up-regulated and between −1.5 to +1.5 are termed as neutrally regulated.

### Gene ontology (GO) and pathway analysis

Over representation analysis for the biological process, Molecular function, Cellular component was performed using ClusterProfiler R Bioconductor package (44). GO related information was obtained from biomaRt (45) R package. Gene Ontology (GO) with multiple test adjusted p-value ≤ 0.05 are considered significant. To visualize the GO enrichment results, GOplot R package was used. GOplot package calculates z-score using the following formula,

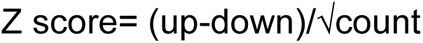

Where “up” is the number of up regulated genes in a GO term and “down” represents number of down regulated genes in the GO term. The z-score provides a rough idea about the expression profile of genes within a GO term (46).

### Quantitative Real Time-PCR and analysis

Total RNA was isolated as described above and was quantified using Nanodrop 2000 spectrophotometer (Thermo Scientific, USA). cDNA synthesis was performed using approximately 1 µg RNA was taken using the Revert Aid H Minus First Strand cDNA Synthesis Kit (Thermo Fisher Scientific, Waltham, MA, United States, Catalog No: K2562) according to the manufacturer’s instructions. iTaq Universal SYBR green supermix (Bio-Rad, Catalog No: 172-5124) was used along with the desired gene-specific oligonucleotide primers (Table S2) to evaluate the quantitative expression profile after normalization with the housekeeping gene *CauACT1* using CFX96TM real-time PCR system (Bio-Rad, USA). The gene expression level was measured by calculating the threshold cycle (C_T_) value of the housekeeping gene, *CauACT1* gene and the desired target genes. Comparative gene expression profiles were measured by the 2^-ΔΔCT^ method. qRT-PCR was performed in biological duplicates and technical triplicates.

### Genomic DNA isolation

Genomic DNA was extracted from the cells grown in YEPD liquid using the Qiagen Yeast DNA Kit (QIAamp DNA Mini Kit, Cat No 51304) according to the manufacturer’s instructions. Genomic DNA was then eluted with distilled water and concentration (absorbance at 260 nm) and purity (ratio absorbance at 260 nm/280 nm) was checked using Nanodrop 2000 spectrophotometer (Thermo Scientific, USA). Whole genomic sequencing was performed at the Clevergene Biocorp Pvt Ltd, Bangalore India.

### Whole genome sequencing and Data analysis

For genome sequencing, a paired-end library with an average insert size of 300 bp was prepared and sequenced using the Illumina NovaSeq 6000 platform. Data quality was checked using FastQC and MultiQC (47). The data was checked for base call quality distribution, % bases above Q20, Q30, %GC, and sequencing adapter contamination. All the samples passed the QC threshold (Q20>95%). Raw sequence reads were processed to remove adapter sequences and low-quality bases using fastp (48).

### Alignment, Variant Calling, and Variant Annotation

The trimmed reads were aligned to the reference genome of *Candida auris* strain B8441 (http://www.candidagenome.org/download/sequence/C_auris_B8441) using the bwa mem algorithm (49). The alignments were processed to remove PCR duplicates using samtools (50). The base qualities were recalibrated, and variants were called using freebayes (https://github.com/freebayes/freebayes) with haplotype 1 and a quality score more than 20. The variants were annotated using snpEff (51) using the annotation of their respective reference genomes. The vcf files were processed using snpSift (52) to convert the data into tab-delimited text format files, and the common variants in all the 3 samples were identified using VCF tools.

### Electrophoretic karyotyping

Overnight cultures derived from single colonies were used as inoculum for secondary cultures, which were grown till OD_600_ =0.9. Approximately 3 OD cells were used for chromosomal plug preparation, following the manufacturer’s instructions (Biorad), using Cleancut agarose (0.6%), lyticase enzyme, and Proteinase K. The chromosomes embedded in the agarose plugs were separated on a 1.0% agarose gel (Biorad) using 0.5X TBE as the running buffer. The run protocol is as follows: 60-60 sec switch, 6V/cm, 120° over 8 hours at 12°C, followed by 90-150 sec switch, 6V/cm, 120° over 18 hours at 12°C. The run was performed in CHEF-DR III system (Biorad). The gel was stained with ethidium bromide post-run, and the bands were visualized using Gel documentation system (Biorad).

### R6G efflux assay

The R6G (Rhodamine 6G) efflux assay was performed by the energy-dependent efflux method. In this assay, *C. auris* cells from an overnight culture were inoculated in fresh YEPD medium at 0.1 OD and grown for 4-5 h at 30°C until the log phase. The cell suspension was washed with 1X PBS (phosphate buffered saline) twice and incubated for 3 h at 200 rpm and 30°C for starvation (glucose-free) to reduce the activity of the ABC transporters. After incubation, cells were washed twice with PBS and diluted to obtain 10^8^ cells/mL in PBS. R6G at a final concentration of 10 µM was added to the suspension and incubated for 3 hrs at 30°C and 200 rpm for accumulation assay. For the efflux assay, the cells were washed twice in PBS, 2% glucose was added to the suspension, and incubated for 45 min. The supernatant was then collected, followed by measurement of fluorescence of R6G in a fluorescence spectrophotometer at excitation and emission wavelengths of 527 nm and 555 nm, respectively.

### Sterol measurements

Sterols were extracted using the alcoholic KOH method and the percentage of ergosterol was calculated as per the below mentioned method:

The extracted sterols indicated a four-peak spectral absorption pattern produced by ergosterol and 24(28)-dehydroergosterol (DHE) contents. Both ergosterol and DHE absorb at 281.5 nm, whereas only DHE absorbs at 230 nm. The ergosterol content is determined by subtracting the amount of DHE (calculated from A_230_ nm) from the total ergosterol plus DHE content (calculated from A_281.5_ nm). The ergosterol content was calculated as a percentage of the wet weight of the cells using the following equations:

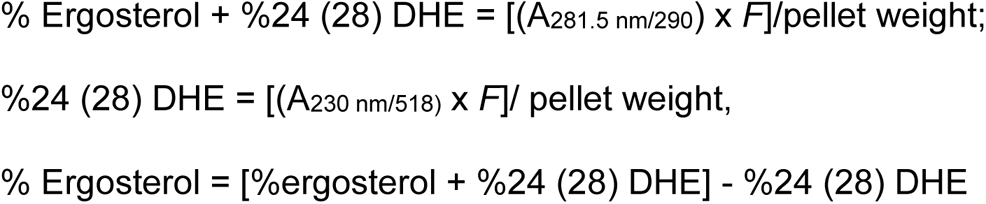

Where *F* is the dilution factor in petroleum ether and 290 and 518 are the E values (in percent per centimeter) determined for crystalline ergosterol and 24(28) DHE, respectively (53).

### ROS estimation

A saturated, overnight culture was used to inoculate 10 mL of fresh YEPD at an OD of 0.1, which was grown for 5-6 hours at 30°C to obtain a log phase culture. The cells were then collected by centrifugation and washed with autoclaved PBS buffer. The cells were then incubated with 10µM DCFDA for 30 minutes in dark. Post incubation, the cells were washed and resuspended in 1 mL PBS for fluorescence measurement. The excitation and emission wavelengths taken were 480 nm and 540 nm respectively.

### Lipid extraction

Saturated overnight cultures in YEPD were diluted to 0.1 OD_600_ in fresh media and grown up to OD_600_ 0.8 to 1 (mid-log phase). In three biological replicates, approximately 5× 10^8^ cells of each strain were harvested by centrifugation at 4000xg for 5 minutes. Pellets were washed twice with sterile water. C17 Sphingosine and C17 Ceramide (d18:1, 17:0) (Avanti Polar Lipids, USA), as internal standards, were added to each sample and cells were lysed with the glass beads (50 mg, 0.4-0.6mm) in Fastprep® (MP Biomedical). Lipid extraction and base hydrolysis was performed using the methods described earlier by (26). Extracted lipids were N_2_ dried and stored at −20°C until analyzed.

### Protein estimation

Protein estimation was done using Bicinchoninic Acid (BCA) Protein Assay kit (G-Biosciences). Cell lysate (25ul aliquot of each replicate) was diluted with ddH_2_O (1:8) in 96-well round bottom plates, and absorbance was read at 595 nm. A standard curve based on serial dilutions of the Bovine Serum Albumin (BSA) as a standard was used for calibration. The amount of protein (mg/mL) was calculated from the slope of the standard calibration curve.

### Liquid Chromatography Mass Spectrometry

Extracted lipids were suspended in a buffer consisting of methanol containing 1mM ammonium formate + 0.2 % formic acid (organic buffer). A two-buffer mobile system was used: Water containing 2 mM ammonium formate + 0.2% formic acid (aqueous buffer) and organic buffer. Autosampler delivered 5 µL sample, and pumps fetched mobile buffer at a flow rate of 0.3 mL per minute to the HPLC fitted with the column. The C8 column (Waters, USA) was used to separate SLs. SL species were detected by multiple reaction monitoring (MRM) methods using QTRAP® 4500 (SCIEX, USA) mass spectrometer.

### Data Analysis and statistical analysis

Mass spectrometric chromatograms were resolved using MultiQuant^TM^ software (SCIEX, USA). Quantification was done using the internal standard normalization method. The data was further normalized to per mg protein, and the amount of each lipid species was calculated as % ng per mg protein. Three replicates of each sample were used for all analyses. Statistical significance between the data sets was determined using Student’s *t*-test, and a *p*-value of≤ 0.05 was considered significant. Data bars were plotted using GraphPad Prism 8.

## Supporting information

Supplemental sheet

## Data Availability

The raw reads of RNA sequencing and whole genome sequencing have been deposited in NCBI database under the Bioproject PRJNA1012821.

## ACKNOWLEDGEMENTS

This study was funded by the Indian Council of Medical Research (Myco/Adhoc/1/2022-ECD-II), the Government of India, to R.P., A.C., S.M.R., and K.S. R.P. acknowledges the departmental grants, DBT Builder Program (BT/INF/22/SP45072/2022), and DBT PG TEACHING (BT/HRD/01/46/2020) awarded to Amity Institute of biotechnology, Amity University, Haryana. A.C. acknowledges the Indian Council of Medical Research, Government of India, for the award of a Senior Research fellowship ((Myco/Fell/7/2022-ECD-II)). P.K. acknowledges the fellowship support received from ICMR (2020-7196/CMB-BMS). M.K. acknowledges the award of ICMR-RA fellowship (Myco/Fell/3/2022-ECD-II). A.N. acknowledges JNCASR for the postdoctoral fellowship. A.C. acknowledges the support of the Amity Central Instrument Research facility (CIRF) and Amity Lipidomics Research Facility (ALRF) in carrying out part of the work.

## AUTHOR CONTRIBUTIONS

AC- Methodology, Investigation, Validation, Data curation, Formal analysis, Visualization, Writing-original draft

PK- Formal analysis

MK- Data curation, Visualization, Formal analysis

AN- Data curation, Investigation, Formal analysis

KY- Data curation, Investigation, Formal analysis

BA- Data curation, Visualization, Formal analysis

AS- Data curation, Formal analysis

AS- Data curation, Formal analysis

AB- Visualization, Formal analysis

SMR- Funding acquisition, Supervision, Resources

AC- Funding acquisition, Supervision, Resources

AKM- Supervision, Resources

NAG- Supervision, Resources

KS- Funding acquisition, Supervision, Resources

RP- Methodology, Conceptualization, Funding acquisition, Project administration, Supervision, Original draft- review & editing

---

**FIG S1:**
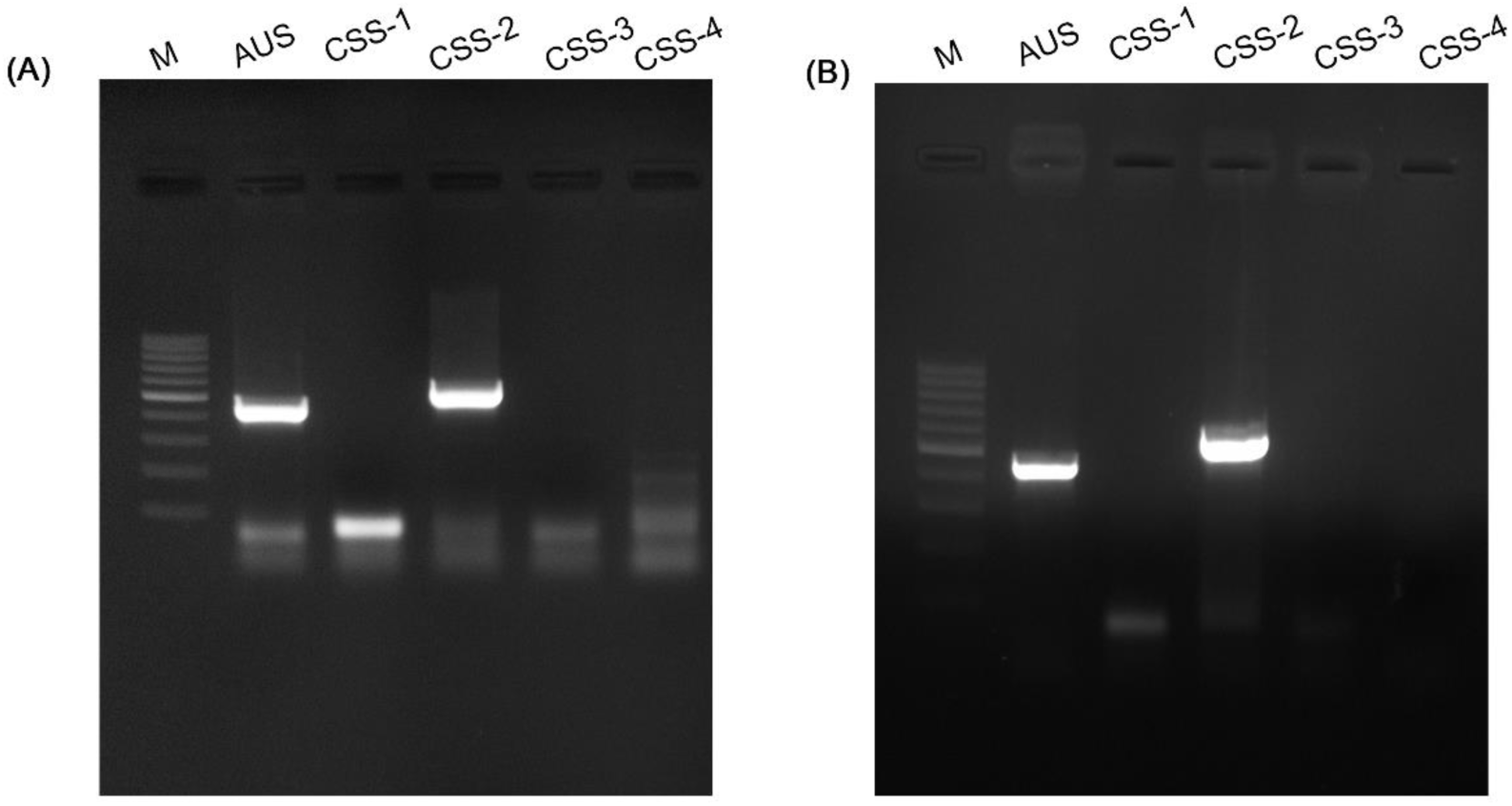
Clade typing for the two isolates used in this study. AUS-Auris universal sequence primers, amplify DNA sequence from all the four clades, CSS-I-amplicon of clade specific sequence for clade I, CSS-II-amplicon of clade specific sequence for clade II, CSS-III-amplicon of clade specific sequence for clade III, and CSS-IV-amplicon of clade specific sequence for clade IV. The molecular size marker (100 bp ladder) is labelled as M. **(A)** Clade-typing of the strain CBS10913^T^, and **(B)** Clade-typing of the strain P2428.

**FIG S2:**
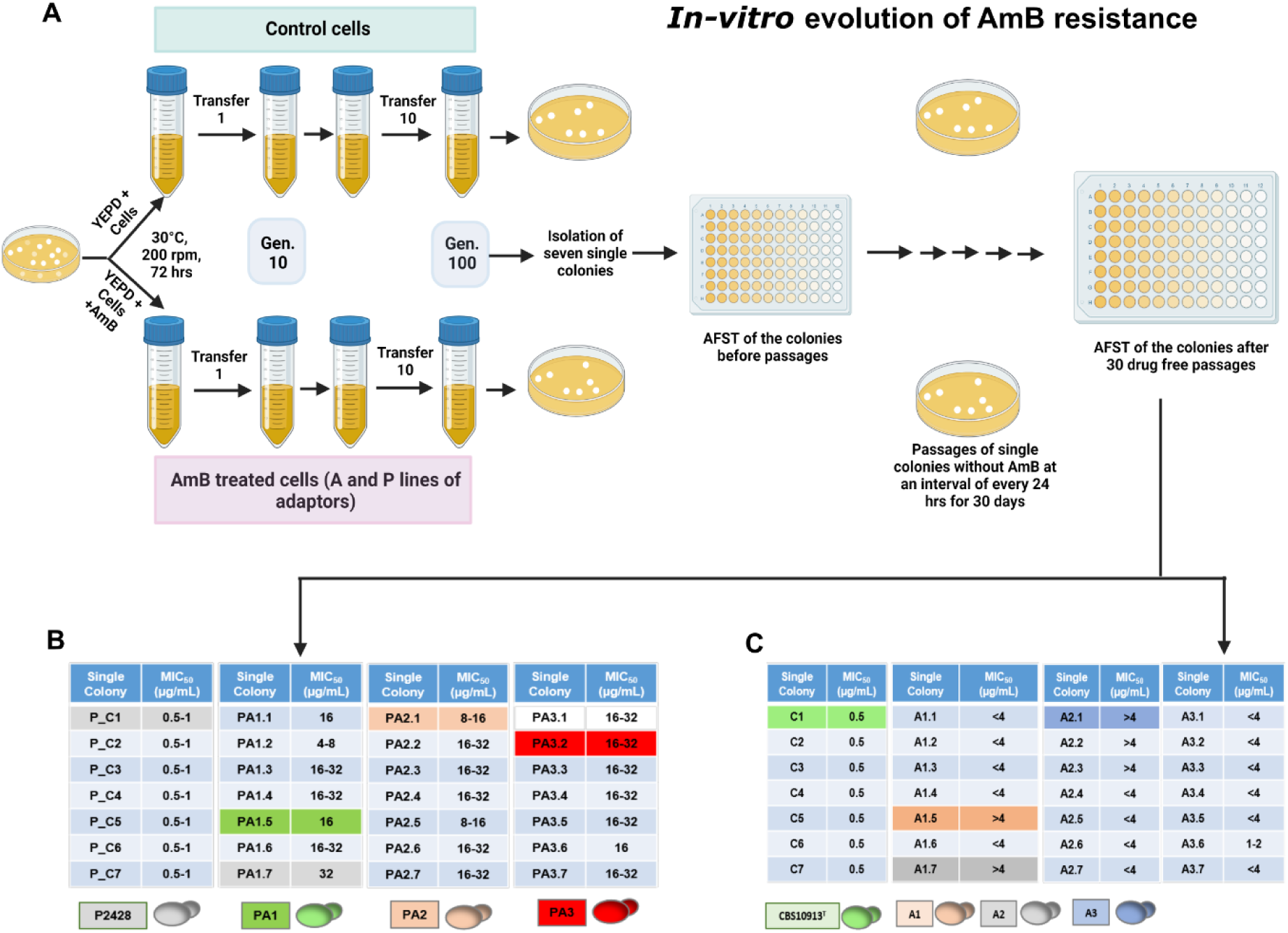
In-vitro evolution methodology and selection of the final colonies retaining the high resistance pst drug free passages. **A.** Schematic depicting the in-vitro microevolution regime followed in the present study followed by **B and C.** Selection of highly resistant single colony for each adapted replicate post adaptation from the isolated single colonies of both the adaptor lines of clade II (P line and A line) after 30 drug free passages done at every 24 hrs interval.

**FIG S3:**
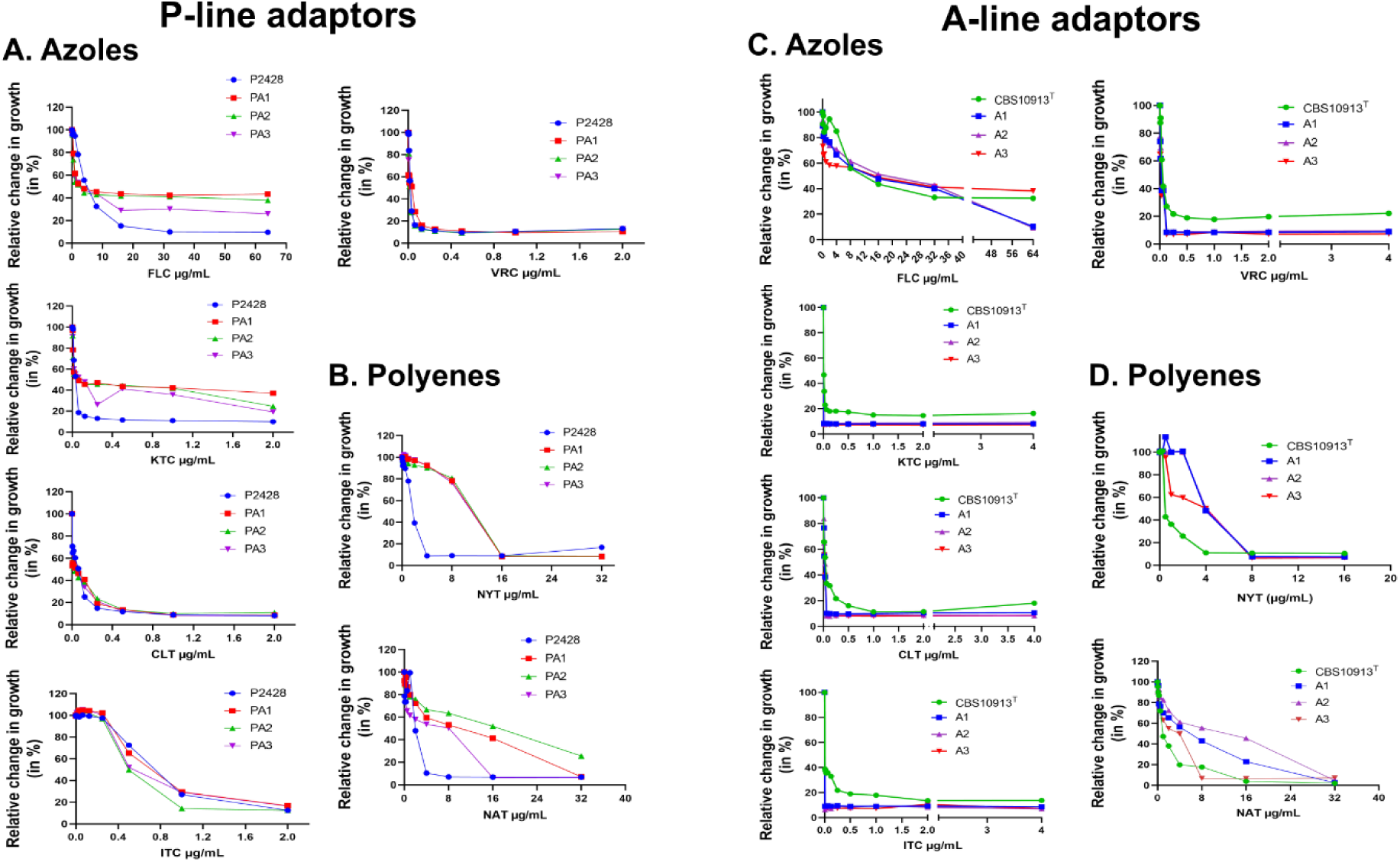
Cross-resistance of P-line (left panel), and A-line (right panel) adaptor strains along with their respective controls. (P2428, and CBS10913^T^) representing growth differences by line plots towards different drugs. **(A, and C)** Fluconazole (FLC), ketoconazole (KTC), clotrimazole (CLT), itraconazole (ITC), voriconazole (VRC). **(B, and D)** Growth after exposure with polyenes, nystatin (NYT), and natamycin (NAT). The line plots depict the drug concentrations on the x-axis, and growth on the y-axis as a relative percentage change in growth by a micro-cultivation method in a 96-well round bottom plate using a multimode reader (Tecan Infinite M Plex, USA) in YEPD broth at 30 °C. All experiments were done in triplicates and standard deviations were calculated

**FIG S4:**
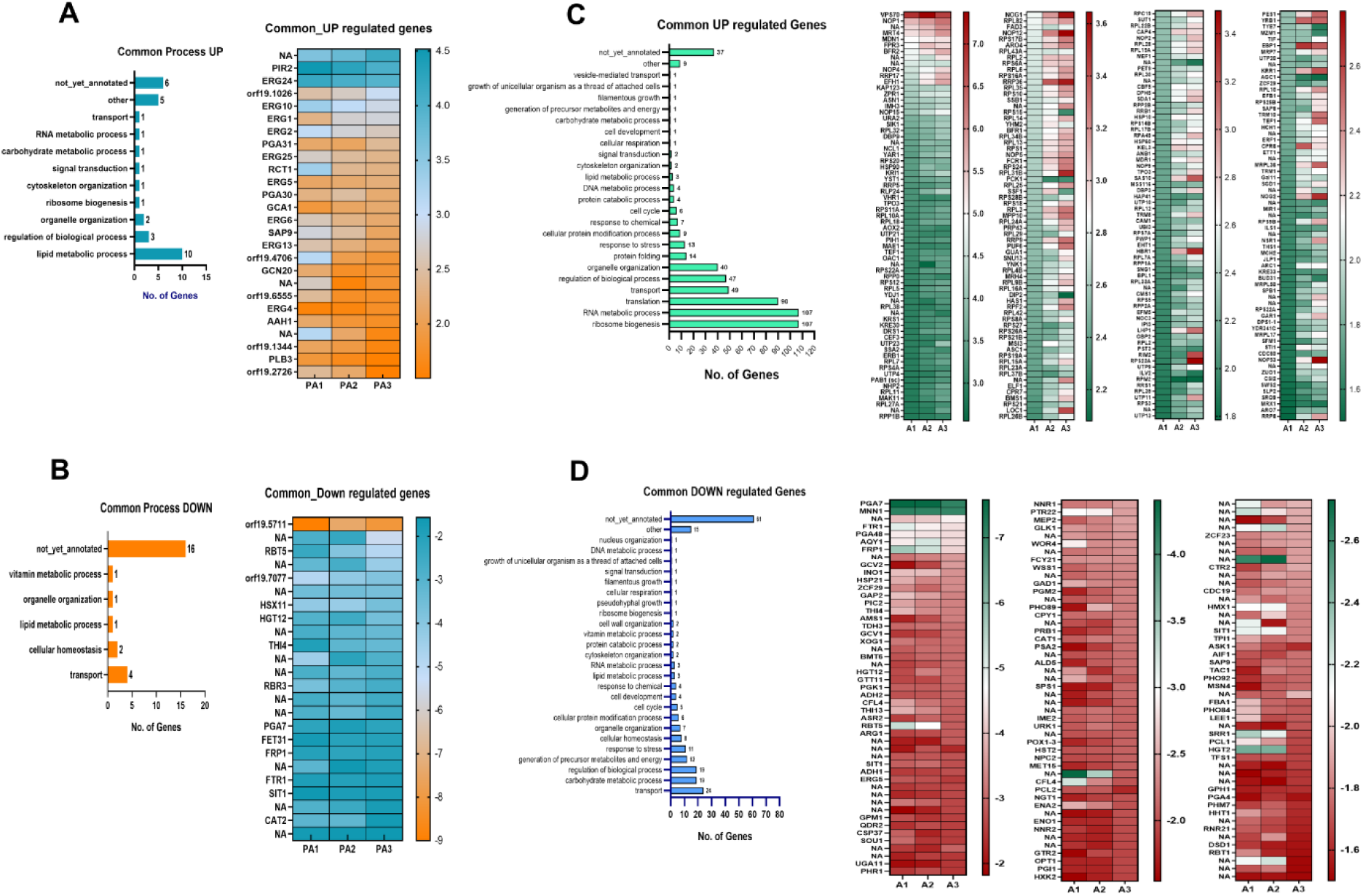
Total number of Common DEGs. The respective Up and Down-regulated genes depicted as heatmaps along with the categories of genes participating in various cellular process depicted by bar graphs in **(A, B)** P-line and, **(C, D)** A-line of adaptors.

**FIG S5:**
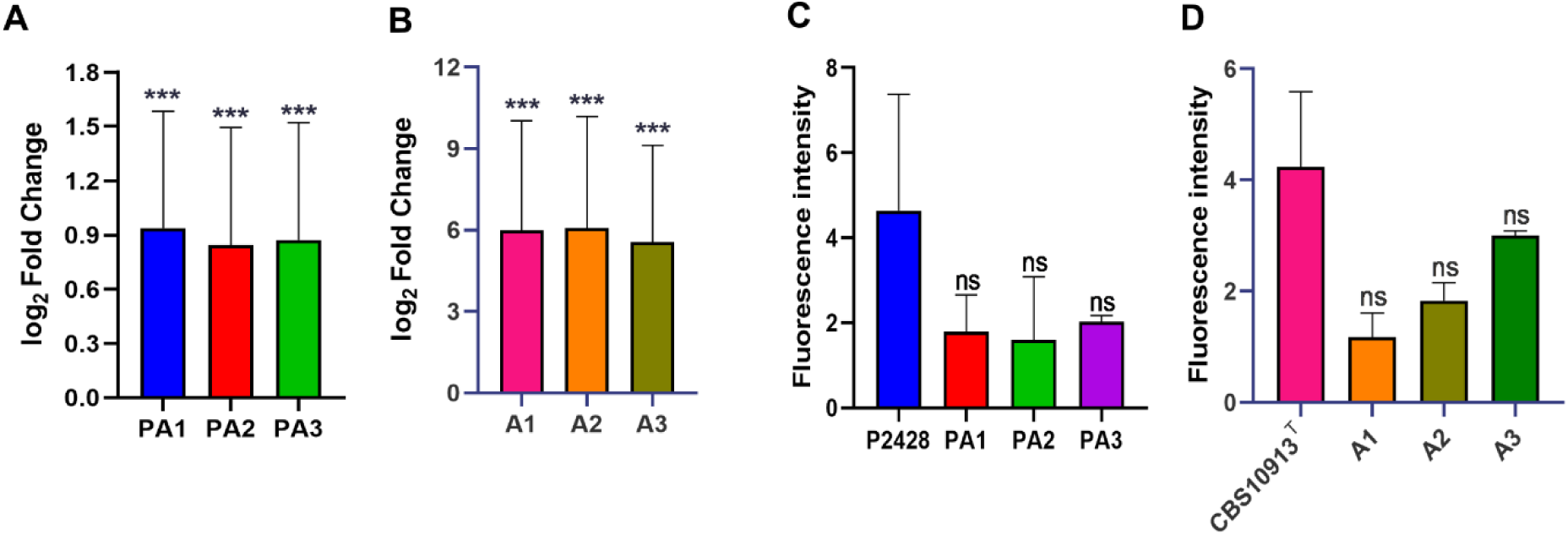
The relationship between of *AOX2* and ROS. **A** The *AOX2* expression levels in the adaptors of both the P, and **B**, A lines. **C, and D** The ROS measurement in the adaptors. With the increase in *AOX2* expression, the ROS levels were decreased.

**FIG S6:**
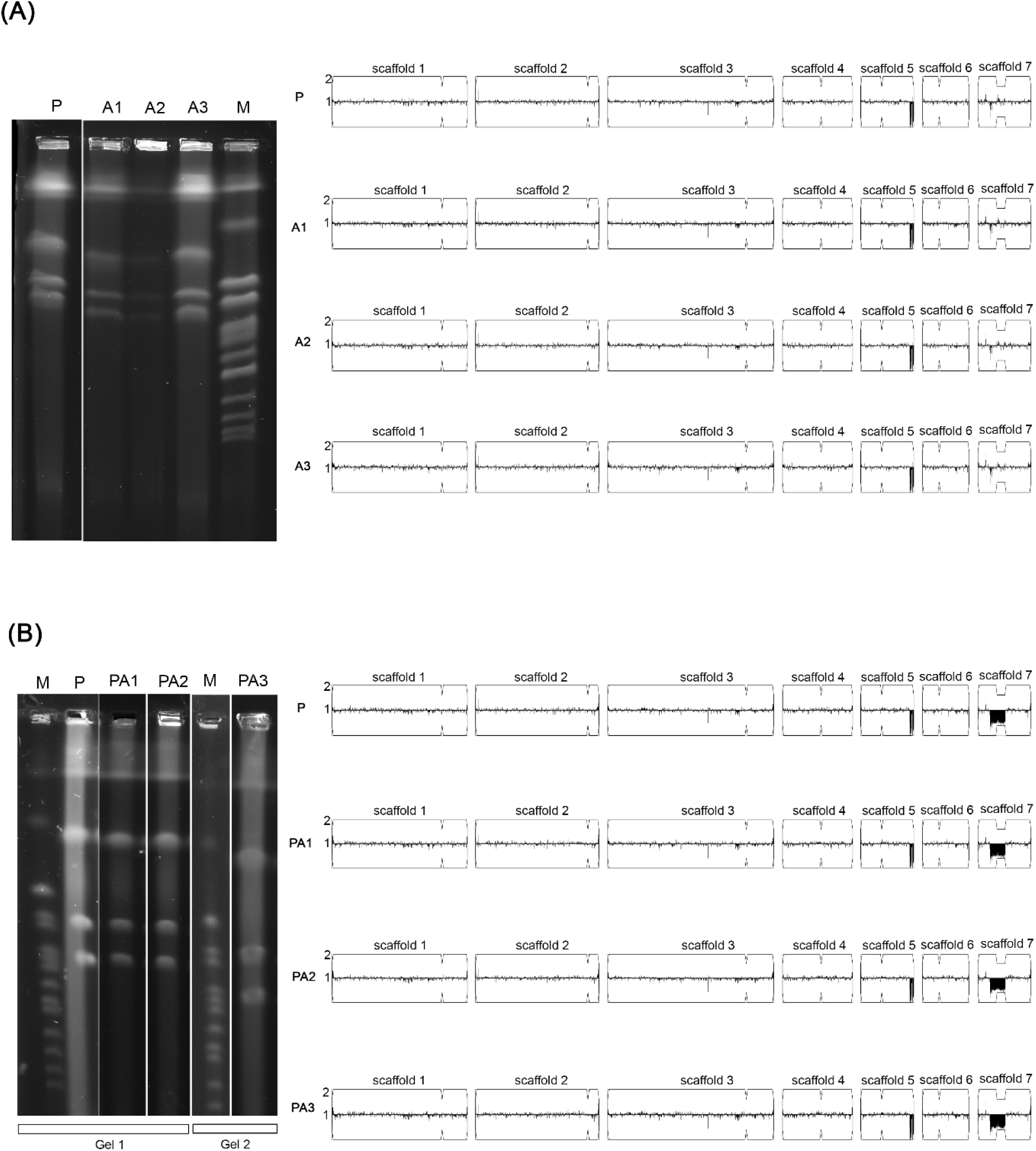
Ploidy changes analysis in the A-, and P-lines of the AmB adaptors. (A) Karyotype analysis and CNV analysis of CBS10913^T^ (Parent, P) and the adaptors A1, A2 and A3. (B) Karyotype analysis and CNV analysis of P2428 (Parent, P) and the adaptors PA1, PA2 and PA3. For karyotype analysis, *S. cerevisiae* chromosomes are used as molecular size markers (M). For the CNV analysis, the scaffold numbers and the centromere locations (breaks) are shown.

**FIG S7:**
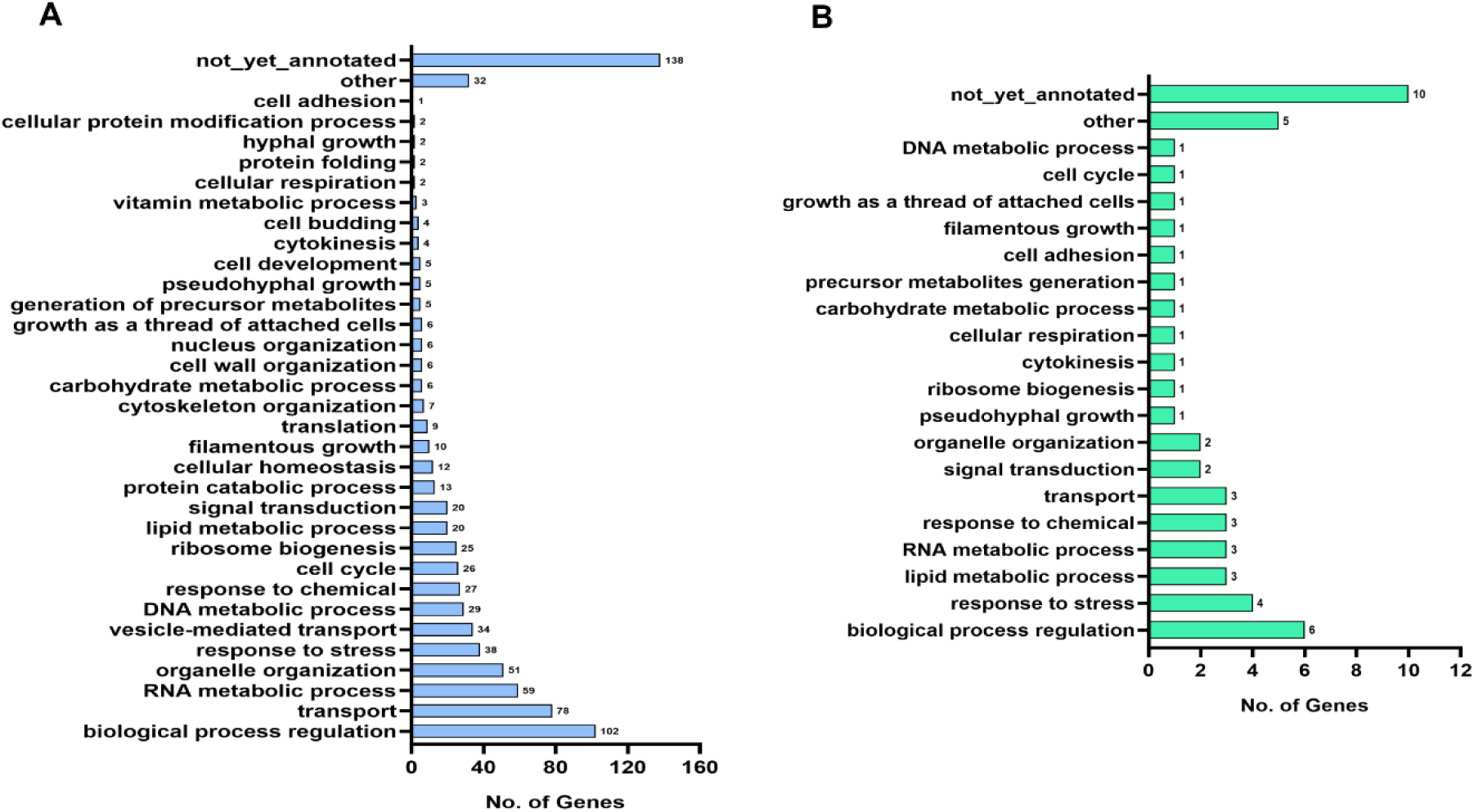
The landscapes of gene expression with SNPs. **(A)** P-line, and **(B)** A-line. The bar graphs depict number of genes related to a particular cell process that are having SNPs. The cell processes were found out by slim mapping of the genes exhibiting SNPs via candida genome database.

