## Supplemental tables for "Microevolution of clade II isolates of *Candida auris* highlights multifaceted intra-clade heterogeneity in acquiring resistance towards amphotericin B"

**Supplementary Tables**

**Table S1: Strains used in the study**

| **Strain** | **Description** | **Source** |
| --- | --- | --- |
| P2428 | Clade I type AmB susceptible strain | National Culture Collection of Pathogenic Fungi (NCCPF), Indian Council of Medical Research (ICMR), New Delhi, sponsored National facility at the Mycology Division, Department of Medical Microbiology, Postgraduate Institute of Medical Education and Research (PIGMER), Chandigarh, India |
| PA1 | Strain obtained from the terminal line of the *in vitro* evolution experiment | This study |
| PA2 | Strain obtained from the terminal line of the *in vitro* evolution experiment | This study |
| PA3 | Strain obtained from the terminal line of the *in vitro* evolution experiment | This study |
| CBS 109A3T | Clade II type AmB susceptible strain | Central Bureau voor Schimmel Cultures (CBS), Fungal Biodiversity Centre of the Royal Netherlands Academy of Arts and Sciences (KNAW), Utrecht |
| A1 | Strain obtained from the terminal line of the *in vitro* evolution experiment | This study |
| A2 | Strain obtained from the terminal line of the *in vitro* evolution experiment | This study |
| A3 | Strain obtained from the terminal line of the *in vitro* evolution experiment | This study |
| AmB^R^ (470140) | Clade I type AmB resistant clinical isolate | National Culture Collection of Pathogenic Fungi (NCCPF), Indian Council of Medical Research (ICMR), New Delhi, sponsored National facility at the Mycology Division, Department of Medical Microbiology, Postgraduate Institute of Medical Education and Research (PIGMER), Chandigarh, India |
| FLC^R^ (470147) | Clade I type FLC resistant clinical isolate | National Culture Collection of Pathogenic Fungi (NCCPF), Indian Council of Medical Research (ICMR), New Delhi, sponsored National facility at the Mycology Division, Department of Medical Microbiology, Postgraduate Institute of Medical Education and Research (PIGMER), Chandigarh, India |

**Table S2:** List of primers used in this study

FP (forward primer), RP (reverse primer), and RT (Real Time)

| **Primer name** | **Primer sequence (5’- - -3’)** |
| --- | --- |
| B9J08_001930 RT FP  B9J08_001930 RT RP | GACTCCTACTCATCGTGTTC  GTTCATCTCCCATCTGGTGC |
| B9J08_003910 RT FP  B9J08_003910 RT RP | CAACAGACAGTTCCAGTTTG  GAAGTCTCCAGAAAGACATTTG |
| B9J08_004469 RT FP  B9J08_004469 RT RP | AAGTGGTGGCTCTGTGTATT  GAAACCGAAGAAGGCATGTT |
| B9J08_000675 RT FP  B9J08_000675 RT RP | CAACTACACCCTAGGATCTTT  CCCGAAGGTATTTACGATTTC |
| B9J08_000438 RT FP  B9J08_000438 RT RP | CTGTAAAGACTTGAACGATCC  AGAACTTGACTCAGAAGATCC |
| B9J08_004476 RT FP  B9J08_004476 RT RP | GAATGACCCATACCACTACTTT  TAATAGAGGCAGATGGCTTTG |
| B9J08_003251 RT FP  B9J08_003251 RT RP | TATATAACGCCGTCTCTCTTC  GTCTTCAGTGTACCATGTATC |
| B9J08_000918 RT FP  B9J08_000918 RT RP | CGAATCCCAGGATAAGTATGT  GTACACTCTCAAGACATTGGT |
| B9J08_005245 RT FP  B9J08_005245 RT RP | GAGCTTGGAATCAACACTATC  CTCTAGCAATAGACGCTTTAG |

**Table S3:** List Non-synonymous mutations present exclusively in PA1

| **Chromosome** | **POS** | **Gene ID** | **REF** | **ALT** |
| --- | --- | --- | --- | --- |
| PEKT02000001 | 174439 | B9J08_000081 | T | C |
| PEKT02000001 | 174442 | B9J08_000081 | G | T |
| PEKT02000001 | 389337 | B9J08_000181 | C | T |
| PEKT02000001 | 389338 | B9J08_000181 | A | T |
| PEKT02000001 | 443241 | B9J08_000204 | G | A |
| PEKT02000001 | 639957 | B9J08_000315 | A | G |
| PEKT02000001 | 639962 | B9J08_000315 | T | C |
| PEKT02000001 | 662236 | B9J08_000325 | A | C |
| PEKT02000001 | 683679 | B9J08_000336 | G | C |
| PEKT02000001 | 690594 | B9J08_000338 | T | C |
| PEKT02000001 | 690603 | B9J08_000338 | A | C |
| PEKT02000001 | 704346 | B9J08_000345 | C | T |
| PEKT02000001 | 814786 | B9J08_000398 | T | G |
| PEKT02000001 | 972383 | B9J08_000458 | G | A |
| PEKT02000002 | 83745 | B9J08_000539 | C | T |
| PEKT02000002 | 138264 | B9J08_000572 | T | A |
| PEKT02000002 | 164822 | B9J08_000583 | G | T |
| PEKT02000002 | 285316 | B9J08_000640 | A | T |
| PEKT02000002 | 301282 | B9J08_000651 | C | A |
| PEKT02000002 | 305778 | B9J08_000653 | T | A |
| PEKT02000002 | 306081 | B9J08_000653 | A | G |
| PEKT02000002 | 392443 | B9J08_000691 | T | G |
| PEKT02000002 | 400034 | B9J08_000693 | G | A |
| PEKT02000002 | 417677 | B9J08_000701 | G | T |
| PEKT02000002 | 432798 | B9J08_000706 | C | T |
| PEKT02000002 | 456542 | B9J08_000717 | A | G |
| PEKT02000002 | 519705 | B9J08_000747 | A | T |
| PEKT02000002 | 679190 | B9J08_000815 | G | A |
| PEKT02000002 | 697699 | B9J08_000824 | C | T |
| PEKT02000002 | 705192 | B9J08_000828 | A | G |
| PEKT02000002 | 755930 | B9J08_000850 | T | C |
| PEKT02000002 | 769640 | B9J08_000857 | C | T |
| PEKT02000002 | 772720 | B9J08_000858 | A | T |
| PEKT02000002 | 898598 | B9J08_000921 | G | A |
| PEKT02000002 | 914889 | B9J08_000926 | C | T |
| PEKT02000002 | 932680 | B9J08_000931 | A | G |
| PEKT02000002 | 932910 | B9J08_000931 | G | T |
| PEKT02000002 | 968186 | B9J08_000943 | A | C |
| PEKT02000002 | 981366 | B9J08_000950 | C | T |
| PEKT02000002 | 989556 | B9J08_000957 | C | T |
| PEKT02000002 | 1043556 | B9J08_000976 | A | C |
| PEKT02000002 | 1051699 | B9J08_000980 | C | A |
| PEKT02000002 | 1075283 | B9J08_000990 | G | A |
| PEKT02000002 | 1194450 | B9J08_001043 | A | G |
| PEKT02000002 | 1217228 | B9J08_001053 | C | T |
| PEKT02000003 | 91327 | B9J08_001107 | G | A |
| PEKT02000003 | 111943 | B9J08_001119 | G | C |
| PEKT02000003 | 172607 | B9J08_001142 | T | C |
| PEKT02000003 | 175845 | B9J08_001144 | A | C |
| PEKT02000003 | 183632 | B9J08_001149 | T | C |
| PEKT02000003 | 282788 | B9J08_001197 | C | T |
| PEKT02000003 | 397829 | B9J08_001254 | G | A |
| PEKT02000003 | 409721 | B9J08_001259 | T | C |
| PEKT02000003 | 530012 | B9J08_001307 | T | G |
| PEKT02000003 | 530553 | B9J08_001307 | A | T |
| PEKT02000003 | 582143 | B9J08_001331 | C | T |
| PEKT02000003 | 602418 | B9J08_001343 | A | T |
| PEKT02000003 | 627592 | B9J08_001355 | T | C |
| PEKT02000003 | 641160 | B9J08_001361 | T | C |
| PEKT02000003 | 706260 | B9J08_001383 | C | T |
| PEKT02000003 | 803667 | B9J08_001431 | C | T |
| PEKT02000003 | 813401 | B9J08_001437 | G | T |
| PEKT02000003 | 902137 | B9J08_001471 | C | T |
| PEKT02000003 | 995485 | B9J08_001511 | C | T |
| PEKT02000004 | 37142 | B9J08_001544 | C | T |
| PEKT02000004 | 40700 | B9J08_001546 | T | A |
| PEKT02000004 | 48002 | B9J08_001550 | G | T |
| PEKT02000004 | 83070 | B9J08_001565 | A | G |
| PEKT02000004 | 212745 | B9J08_001624 | T | C |
| PEKT02000004 | 219078 | B9J08_001626 | T | A |
| PEKT02000004 | 229681 | B9J08_001634 | A | G |
| PEKT02000004 | 237787 | B9J08_001641 | G | A |
| PEKT02000004 | 384658 | B9J08_001707 | A | T |
| PEKT02000004 | 404952 | B9J08_001717 | G | C |
| PEKT02000004 | 412412 | B9J08_001721 | A | T |
| PEKT02000004 | 495547 | B9J08_001758 | T | A |
| PEKT02000004 | 499133 | B9J08_001760 | G | A |
| PEKT02000004 | 501091 | B9J08_001761 | C | T |
| PEKT02000004 | 597472 | B9J08_001808 | C | T |
| PEKT02000004 | 694433 | B9J08_001857 | T | C |
| PEKT02000004 | 763854 | B9J08_001898 | C | G |
| PEKT02000004 | 780842 | B9J08_001902 | G | A |
| PEKT02000005 | 34530 | B9J08_001957 | G | A |
| PEKT02000005 | 58324 | B9J08_001968 | T | C |
| PEKT02000005 | 76011 | B9J08_001975 | A | T |
| PEKT02000005 | 76743 | B9J08_001975 | C | A |
| PEKT02000005 | 100387 | B9J08_001988 | T | A |
| PEKT02000005 | 387440 | B9J08_002125 | G | A |
| PEKT02000005 | 403286 | B9J08_002131 | C | G |
| PEKT02000005 | 424105 | B9J08_002139 | T | C |
| PEKT02000005 | 435334 | B9J08_002144 | T | C |
| PEKT02000005 | 440873 | B9J08_002146 | C | T |
| PEKT02000005 | 471194 | B9J08_002161 | T | A |
| PEKT02000005 | 518352 | B9J08_002181 | T | C |
| PEKT02000005 | 547327 | B9J08_002193 | T | G |
| PEKT02000005 | 579191 | B9J08_002209 | A | T |
| PEKT02000005 | 581951 | B9J08_002211 | A | G |
| PEKT02000006 | 21146 | B9J08_002244 | C | A |
| PEKT02000006 | 115243 | B9J08_002283 | A | G |
| PEKT02000006 | 119733 | B9J08_002286 | G | A |
| PEKT02000006 | 195647 | B9J08_002318 | G | A |
| PEKT02000006 | 197428 | B9J08_002319 | C | T |
| PEKT02000006 | 214470 | B9J08_002325 | C | G |
| PEKT02000006 | 279609 | B9J08_002354 | G | A |
| PEKT02000006 | 334853 | B9J08_002383 | A | G |
| PEKT02000006 | 376561 | B9J08_002405 | A | G |
| PEKT02000006 | 378005 | B9J08_002407 | A | C |
| PEKT02000006 | 519159 | B9J08_002465 | G | A |
| PEKT02000006 | 534553 | B9J08_002471 | T | G |
| PEKT02000006 | 578192 | B9J08_002489 | G | A |
| PEKT02000006 | 590268 | B9J08_002494 | C | T |
| PEKT02000006 | 634765 | B9J08_002518 | G | T |
| PEKT02000006 | 687775 | B9J08_002539 | C | G |
| PEKT02000006 | 698362 | B9J08_002544 | A | T |
| PEKT02000006 | 754464 | B9J08_002575 | T | C |
| PEKT02000006 | 757506 | B9J08_002577 | G | A |
| PEKT02000007 | 75495 | B9J08_002620 | G | C |
| PEKT02000007 | 103808 | B9J08_002634 | A | C |
| PEKT02000007 | 104029 | B9J08_002634 | G | T |
| PEKT02000007 | 109013 | B9J08_002636 | C | G |
| PEKT02000007 | 162731 | B9J08_002661 | T | A |
| PEKT02000007 | 170609 | B9J08_002664 | T | A |
| PEKT02000007 | 187300 | B9J08_002667 | T | C |
| PEKT02000007 | 206744 | B9J08_002673 | T | C |
| PEKT02000007 | 259921 | B9J08_002697 | C | T |
| PEKT02000007 | 263163 | B9J08_002698 | C | T |
| PEKT02000007 | 283357 | B9J08_002708 | A | T |
| PEKT02000007 | 285049 | B9J08_002710 | G | A |
| PEKT02000007 | 326507 | B9J08_002734 | G | A |
| PEKT02000007 | 339075 | B9J08_002738 | C | T |
| PEKT02000007 | 394186 | B9J08_002766 | G | A |
| PEKT02000007 | 730753 | B9J08_002931 | G | A |
| PEKT02000007 | 748743 | B9J08_002941 | A | T |
| PEKT02000007 | 813619 | B9J08_002974 | C | A |
| PEKT02000007 | 813850 | B9J08_002974 | C | T |
| PEKT02000007 | 869012 | B9J08_003003 | G | C |
| PEKT02000007 | 881200 | B9J08_003008 | T | A |
| PEKT02000007 | 931920 | B9J08_003040 | A | G |
| PEKT02000007 | 1047985 | B9J08_003106 | A | T |
| PEKT02000007 | 1124530 | B9J08_003147 | G | A |
| PEKT02000007 | 1127589 | B9J08_003150 | C | T |
| PEKT02000007 | 1289437 | B9J08_003229 | G | C |
| PEKT02000007 | 1326425 | B9J08_003242 | C | T |
| PEKT02000007 | 1357094 | B9J08_003259 | G | A |
| PEKT02000007 | 1517777 | B9J08_003333 | A | G |
| PEKT02000007 | 1590043 | B9J08_003373 | C | A |
| PEKT02000007 | 1666044 | B9J08_003406 | C | T |
| PEKT02000007 | 1698545 | B9J08_003421 | A | G |
| PEKT02000007 | 1790942 | B9J08_003460 | G | A |
| PEKT02000007 | 1837906 | B9J08_003482 | T | A |
| PEKT02000007 | 1867698 | B9J08_003498 | T | C |
| PEKT02000007 | 1868359 | B9J08_003499 | C | A |
| PEKT02000007 | 1918045 | B9J08_003522 | G | C |
| PEKT02000007 | 1947447 | B9J08_003541 | A | T |
| PEKT02000007 | 1947577 | B9J08_003541 | A | G |
| PEKT02000007 | 1990090 | B9J08_003562 | G | A |
| PEKT02000007 | 2055078 | B9J08_003589 | T | A |
| PEKT02000007 | 2067171 | B9J08_003591 | A | G |
| PEKT02000007 | 2099886 | B9J08_003609 | C | G |
| PEKT02000007 | 2232979 | B9J08_003669 | T | C |
| PEKT02000007 | 2237481 | B9J08_003672 | G | A |
| PEKT02000007 | 2270406 | B9J08_003694 | T | C |
| PEKT02000007 | 2402331 | B9J08_003754 | A | G |
| PEKT02000007 | 2413339 | B9J08_003761 | T | G |
| PEKT02000007 | 2449907 | B9J08_003775 | G | A |
| PEKT02000007 | 2614481 | B9J08_003845 | T | A |
| PEKT02000007 | 2730374 | B9J08_003896 | C | A |
| PEKT02000007 | 2740751 | B9J08_003900 | A | G |
| PEKT02000007 | 2774533 | B9J08_003915 | A | G |
| PEKT02000007 | 2823072 | B9J08_003936 | G | C |
| PEKT02000007 | 2828053 | B9J08_003937 | A | C |
| PEKT02000007 | 2873796 | B9J08_003956 | A | G |
| PEKT02000007 | 2891347 | B9J08_003965 | G | A |
| PEKT02000007 | 2970424 | B9J08_004003 | A | G |
| PEKT02000007 | 2978613 | B9J08_004009 | C | A |
| PEKT02000007 | 3068434 | B9J08_004046 | C | T |
| PEKT02000007 | 3112404 | B9J08_004071 | G | C |
| PEKT02000008 | 65233 | B9J08_004119 | C | G |
| PEKT02000008 | 129781 | B9J08_004140 | A | T |
| PEKT02000008 | 181340 | B9J08_004159 | A | T |
| PEKT02000008 | 194000 | B9J08_004165 | G | A |
| PEKT02000008 | 215704 | B9J08_004173 | C | G |
| PEKT02000008 | 303035 | B9J08_004210 | T | A |
| PEKT02000008 | 304043 | B9J08_004212 | C | G |
| PEKT02000008 | 307242 | B9J08_004213 | T | G |
| PEKT02000008 | 343495 | B9J08_004232 | C | G |
| PEKT02000008 | 345313 | B9J08_004232 | C | A |
| PEKT02000008 | 388202 | B9J08_004249 | G | A |
| PEKT02000008 | 388353 | B9J08_004249 | T | C |
| PEKT02000008 | 474836 | B9J08_004287 | A | G |
| PEKT02000008 | 531702 | B9J08_004310 | C | T |
| PEKT02000008 | 561368 | B9J08_004322 | A | T |
| PEKT02000008 | 563053 | B9J08_004323 | G | A |
| PEKT02000008 | 565569 | B9J08_004324 | A | G |
| PEKT02000008 | 610671 | B9J08_004344 | T | A |
| PEKT02000008 | 626120 | B9J08_004352 | C | T |
| PEKT02000008 | 691231 | B9J08_004382 | T | G |
| PEKT02000008 | 743859 | B9J08_004401 | A | G |
| PEKT02000008 | 746445 | B9J08_004404 | A | G |
| PEKT02000008 | 766164 | B9J08_004416 | G | A |
| PEKT02000008 | 791170 | B9J08_004428 | G | C |
| PEKT02000008 | 807228 | B9J08_004435 | A | T |
| PEKT02000008 | 811657 | B9J08_004437 | G | C |
| PEKT02000008 | 814953 | B9J08_004438 | C | T |
| PEKT02000008 | 887239 | B9J08_004465 | C | T |
| PEKT02000008 | 888497 | B9J08_004466 | C | G |
| PEKT02000009 | 41190 | B9J08_004481 | A | G |
| PEKT02000009 | 107824 | B9J08_004513 | G | A |
| PEKT02000009 | 111614 | B9J08_004515 | C | T |
| PEKT02000009 | 111656 | B9J08_004515 | C | T |
| PEKT02000009 | 252370 | B9J08_004570 | G | C |
| PEKT02000009 | 317757 | B9J08_004597 | A | G |
| PEKT02000009 | 347729 | B9J08_004611 | A | T |
| PEKT02000009 | 401669 | B9J08_004634 | C | A |
| PEKT02000009 | 491387 | B9J08_004674 | C | T |
| PEKT02000009 | 500081 | B9J08_004677 | G | C |
| PEKT02000009 | 598611 | B9J08_004726 | A | G |
| PEKT02000009 | 612799 | B9J08_004731 | A | T |
| PEKT02000009 | 700386 | B9J08_004773 | C | A |
| PEKT02000009 | 700526 | B9J08_004773 | C | G |
| PEKT02000009 | 710485 | B9J08_004778 | G | C |
| PEKT02000009 | 710800 | B9J08_004778 | T | C |
| PEKT02000009 | 808381 | B9J08_004818 | A | G |
| PEKT02000009 | 893522 | B9J08_004856 | A | G |
| PEKT02000009 | 901256 | B9J08_004860 | C | G |
| PEKT02000009 | 932754 | B9J08_004872 | A | G |
| PEKT02000010 | 97771 | B9J08_004936 | C | A |
| PEKT02000010 | 120651 | B9J08_004948 | C | A |
| PEKT02000010 | 122051 | B9J08_004948 | T | G |
| PEKT02000010 | 177024 | B9J08_004972 | A | G |
| PEKT02000010 | 197322 | B9J08_004985 | A | G |
| PEKT02000010 | 236610 | B9J08_005003 | C | T |
| PEKT02000010 | 248998 | B9J08_005009 | G | T |
| PEKT02000010 | 331159 | B9J08_005049 | C | G |
| PEKT02000010 | 332136 | B9J08_005049 | G | T |
| PEKT02000010 | 336601 | B9J08_005051 | C | T |
| PEKT02000010 | 337098 | B9J08_005051 | G | A |
| PEKT02000010 | 339143 | B9J08_005052 | A | G |
| PEKT02000010 | 339451 | B9J08_005052 | C | T |
| PEKT02000010 | 349764 | B9J08_005057 | C | T |
| PEKT02000010 | 358165 | B9J08_005060 | G | A |
| PEKT02000010 | 411207 | B9J08_005084 | T | A |
| PEKT02000010 | 482442 | B9J08_005117 | A | G |
| PEKT02000010 | 490017 | B9J08_005121 | T | C |
| PEKT02000010 | 499430 | B9J08_005124 | G | C |
| PEKT02000010 | 531577 | B9J08_005143 | G | A |
| PEKT02000010 | 533833 | B9J08_005145 | G | T |
| PEKT02000010 | 534679 | B9J08_005146 | C | G |
| PEKT02000010 | 557034 | B9J08_005158 | G | T |
| PEKT02000010 | 622359 | B9J08_005190 | G | A |
| PEKT02000010 | 628141 | B9J08_005193 | T | C |
| PEKT02000010 | 732370 | B9J08_005234 | C | A |
| PEKT02000010 | 837872 | B9J08_005282 | C | T |
| PEKT02000010 | 898027 | B9J08_005312 | C | T |
| PEKT02000010 | 898876 | B9J08_005313 | T | A |
| PEKT02000010 | 903062 | B9J08_005315 | A | G |
| PEKT02000010 | 912025 | B9J08_005317 | C | G |
| PEKT02000010 | 914087 | B9J08_005318 | T | C |
| PEKT02000010 | 1055616 | B9J08_005382 | G | T |
| PEKT02000010 | 1176211 | B9J08_005428 | C | G |
| PEKT02000010 | 1189512 | B9J08_005430 | G | A |
| PEKT02000010 | 1254986 | B9J08_005464 | C | T |
| PEKT02000010 | 1277905 | B9J08_005478 | T | C |
| PEKT02000010 | 1320275 | B9J08_005500 | G | A |
| PEKT02000010 | 1328108 | B9J08_005504 | A | T |
| PEKT02000010 | 1367955 | B9J08_005521 | A | G |
| PEKT02000010 | 1367994 | B9J08_005521 | T | G |
| PEKT02000010 | 1368619 | B9J08_005521 | T | C |
| PEKT02000010 | 1368739 | B9J08_005521 | A | G |
| PEKT02000010 | 1372554 | B9J08_005522 | A | T |
| PEKT02000010 | 1378300 | B9J08_005524 | G | A |
| PEKT02000012 | 2123 | B9J08_005545 | A | T |
| PEKT02000012 | 5147 | B9J08_005546 | G | C |
| PEKT02000012 | 6684 | B9J08_005547 | G | T |
| PEKT02000012 | 10541 | B9J08_005550 | C | T |
| PEKT02000013 | 3381 | B9J08_005567 | C | T |
| PEKT02000013 | 9653 | B9J08_005569 | T | C |
| PEKT02000013 | 9940 | B9J08_005569 | T | G |
| PEKT02000013 | 10709 | B9J08_005570 | G | T |
| PEKT02000013 | 17006 | B9J08_005572 | T | C |
| PEKT02000014 | 3139 | B9J08_005579 | A | C |
| PEKT02000014 | 3151 | B9J08_005579 | A | C |
| PEKT02000014 | 3174 | B9J08_005579 | G | C |
| PEKT02000014 | 7574 | B9J08_005581 | C | T |
| PEKT02000014 | 8435 | B9J08_005581 | C | T |
| PEKT02000014 | 9021 | B9J08_005581 | C | A |
| PEKT02000014 | 9419 | B9J08_005581 | C | T |
| PEKT02000014 | 9978 | B9J08_005581 | G | T |
| PEKT02000014 | 10357 | B9J08_005581 | C | T |
| PEKT02000014 | 10967 | B9J08_005581 | T | C |

**Table S4:** List Non-synonymous mutations present exclusively in PA2

| **Chromosome** | **POS** | **Gene ID** | **REF** | **ALT** |
| --- | --- | --- | --- | --- |
| PEKT02000001 | 43775 | B9J08_000017 | A | C |
| PEKT02000001 | 132459 | B9J08_000060 | C | G |
| PEKT02000001 | 282360 | B9J08_000131 | A | T |
| PEKT02000001 | 337986 | B9J08_000159 | G | A |
| PEKT02000001 | 344876 | B9J08_000161 | A | C |
| PEKT02000001 | 389337 | B9J08_000181 | C | T |
| PEKT02000001 | 389338 | B9J08_000181 | A | T |
| PEKT02000001 | 443241 | B9J08_000204 | G | A |
| PEKT02000001 | 485584 | B9J08_000228 | T | G |
| PEKT02000001 | 578373 | B9J08_000282 | A | C |
| PEKT02000001 | 639957 | B9J08_000315 | A | G |
| PEKT02000001 | 639962 | B9J08_000315 | T | C |
| PEKT02000001 | 666638 | B9J08_000327 | T | G |
| PEKT02000001 | 683679 | B9J08_000336 | G | C |
| PEKT02000001 | 684960 | B9J08_000336 | A | G |
| PEKT02000001 | 690603 | B9J08_000338 | A | C |
| PEKT02000001 | 704346 | B9J08_000345 | C | T |
| PEKT02000001 | 920410 | B9J08_000440 | T | G |
| PEKT02000001 | 921000 | B9J08_000440 | T | A |
| PEKT02000001 | 972383 | B9J08_000458 | G | A |
| PEKT02000002 | 63952 | B9J08_000526 | A | G |
| PEKT02000002 | 63965 | B9J08_000526 | C | A |
| PEKT02000002 | 227889 | B9J08_000611 | G | T |
| PEKT02000002 | 325409 | B9J08_000661 | A | C |
| PEKT02000002 | 355970 | B9J08_000675 | A | G |
| PEKT02000002 | 455561 | B9J08_000717 | T | A |
| PEKT02000002 | 472864 | B9J08_000725 | G | C |
| PEKT02000002 | 796244 | B9J08_000872 | A | C |
| PEKT02000002 | 840573 | B9J08_000894 | G | A |
| PEKT02000002 | 991428 | B9J08_000958 | G | A |
| PEKT02000002 | 1015759 | B9J08_000966 | A | C |
| PEKT02000002 | 1100549 | B9J08_001002 | T | G |
| PEKT02000003 | 187700 | B9J08_001151 | A | G |
| PEKT02000003 | 193571 | B9J08_001155 | T | C |
| PEKT02000003 | 194135 | B9J08_001155 | G | T |
| PEKT02000003 | 316785 | B9J08_001214 | A | T |
| PEKT02000003 | 316789 | B9J08_001214 | C | A |
| PEKT02000003 | 638068 | B9J08_001359 | A | G |
| PEKT02000003 | 666614 | B9J08_001370 | A | T |
| PEKT02000003 | 738557 | B9J08_001404 | C | T |
| PEKT02000003 | 825765 | B9J08_001443 | C | T |
| PEKT02000003 | 917590 | B9J08_001479 | G | C |
| PEKT02000003 | 985662 | B9J08_001507 | G | A |
| PEKT02000004 | 32807 | B9J08_001542 | G | T |
| PEKT02000004 | 114851 | B9J08_001578 | C | T |
| PEKT02000004 | 165954 | B9J08_001602 | A | C |
| PEKT02000004 | 213182 | B9J08_001624 | A | C |
| PEKT02000004 | 601993 | B9J08_001810 | C | G |
| PEKT02000004 | 602121 | B9J08_001810 | A | C |
| PEKT02000004 | 690654 | B9J08_001855 | T | G |
| PEKT02000004 | 734127 | B9J08_001882 | T | C |
| PEKT02000005 | 97587 | B9J08_001987 | T | G |
| PEKT02000005 | 485116 | B9J08_002169 | T | A |
| PEKT02000005 | 599795 | B9J08_002219 | T | G |
| PEKT02000005 | 636353 | B9J08_002235 | A | G |
| PEKT02000005 | 636358 | B9J08_002235 | G | C |
| PEKT02000006 | 21146 | B9J08_002244 | C | A |
| PEKT02000006 | 37171 | B9J08_002250 | A | C |
| PEKT02000006 | 187702 | B9J08_002313 | A | C |
| PEKT02000006 | 238810 | B9J08_002340 | A | C |
| PEKT02000006 | 238811 | B9J08_002340 | A | C |
| PEKT02000006 | 318905 | B9J08_002374 | A | C |
| PEKT02000006 | 355035 | B9J08_002394 | A | C |
| PEKT02000007 | 53130 | B9J08_002613 | T | C |
| PEKT02000007 | 53131 | B9J08_002613 | G | C |
| PEKT02000007 | 62772 | B9J08_002615 | C | T |
| PEKT02000007 | 231206 | B9J08_002682 | A | G |
| PEKT02000007 | 414153 | B9J08_002775 | A | C |
| PEKT02000007 | 714039 | B9J08_002925 | C | T |
| PEKT02000007 | 763660 | B9J08_002949 | G | C |
| PEKT02000007 | 965421 | B9J08_003057 | T | G |
| PEKT02000007 | 1471164 | B9J08_003310 | C | T |
| PEKT02000007 | 1479829 | B9J08_003314 | A | C |
| PEKT02000007 | 1850770 | B9J08_003489 | T | A |
| PEKT02000007 | 1893488 | B9J08_003509 | C | A |
| PEKT02000007 | 1932061 | B9J08_003531 | T | G |
| PEKT02000007 | 2007198 | B9J08_003571 | A | G |
| PEKT02000007 | 2125240 | B9J08_003619 | T | C |
| PEKT02000007 | 2182731 | B9J08_003643 | T | G |
| PEKT02000007 | 2214997 | B9J08_003661 | A | G |
| PEKT02000007 | 2252372 | B9J08_003683 | C | G |
| PEKT02000007 | 2279404 | B9J08_003698 | G | A |
| PEKT02000007 | 2348817 | B9J08_003727 | C | G |
| PEKT02000007 | 2396511 | B9J08_003751 | G | C |
| PEKT02000007 | 2731321 | B9J08_003896 | G | A |
| PEKT02000007 | 2827767 | B9J08_003937 | T | A |
| PEKT02000007 | 2870704 | B9J08_003955 | A | C |
| PEKT02000007 | 2870715 | B9J08_003955 | C | G |
| PEKT02000007 | 2996163 | B9J08_004015 | T | C |
| PEKT02000007 | 3049345 | B9J08_004036 | G | A |
| PEKT02000007 | 3081123 | B9J08_004053 | A | T |
| PEKT02000007 | 3081124 | B9J08_004053 | C | G |
| PEKT02000007 | 3168758 | B9J08_004097 | A | T |
| PEKT02000008 | 86467 | B9J08_004131 | A | T |
| PEKT02000008 | 216632 | B9J08_004173 | T | G |
| PEKT02000008 | 224267 | B9J08_004176 | T | G |
| PEKT02000008 | 436981 | B9J08_004270 | C | G |
| PEKT02000008 | 622710 | B9J08_004349 | C | G |
| PEKT02000008 | 691029 | B9J08_004382 | T | G |
| PEKT02000008 | 717211 | B9J08_004390 | C | T |
| PEKT02000008 | 779613 | B9J08_004423 | T | G |
| PEKT02000008 | 859650 | B9J08_004453 | T | C |
| PEKT02000008 | 860672 | B9J08_004453 | C | T |
| PEKT02000008 | 887616 | B9J08_004466 | T | C |
| PEKT02000008 | 896291 | B9J08_004469 | G | A |
| PEKT02000009 | 21748 | B9J08_004474 | C | A |
| PEKT02000009 | 21749 | B9J08_004474 | G | C |
| PEKT02000009 | 24589 | B9J08_004475 | A | G |
| PEKT02000009 | 381265 | B9J08_004626 | T | G |
| PEKT02000009 | 383775 | B9J08_004627 | T | G |
| PEKT02000009 | 394518 | B9J08_004631 | C | G |
| PEKT02000009 | 428759 | B9J08_004644 | T | G |
| PEKT02000009 | 880910 | B9J08_004852 | T | G |
| PEKT02000010 | 9941 | B9J08_004898 | T | C |
| PEKT02000010 | 151254 | B9J08_004962 | A | C |
| PEKT02000010 | 174703 | B9J08_004971 | T | C |
| PEKT02000010 | 486023 | B9J08_005119 | T | C |
| PEKT02000010 | 495734 | B9J08_005123 | T | G |
| PEKT02000010 | 513047 | B9J08_005131 | T | A |
| PEKT02000010 | 533833 | B9J08_005145 | G | T |
| PEKT02000010 | 588165 | B9J08_005173 | A | G |
| PEKT02000010 | 588985 | B9J08_005173 | G | T |
| PEKT02000010 | 623757 | B9J08_005191 | T | G |
| PEKT02000010 | 786669 | B9J08_005260 | A | T |
| PEKT02000010 | 786672 | B9J08_005260 | A | T |
| PEKT02000010 | 814666 | B9J08_005272 | C | T |
| PEKT02000010 | 960587 | B9J08_005340 | C | G |
| PEKT02000010 | 1020085 | B9J08_005365 | T | A |
| PEKT02000010 | 1096732 | B9J08_005396 | G | A |
| PEKT02000010 | 1131990 | B9J08_005410 | A | C |
| PEKT02000010 | 1162509 | B9J08_005423 | G | A |
| PEKT02000012 | 2609 | B9J08_005545 | T | G |
| PEKT02000012 | 11113 | B9J08_005550 | T | A |
| PEKT02000012 | 11724 | B9J08_005550 | A | G |
| PEKT02000012 | 11882 | B9J08_005550 | T | A |
| PEKT02000012 | 17957 | B9J08_005551 | C | G |
| PEKT02000012 | 18847 | B9J08_005551 | A | C |
| PEKT02000014 | 3174 | B9J08_005579 | G | C |

**Table S5:** List Non-synonymous mutations present exclusively in PA3

| **Chromosome** | **POS** | **Gene ID** | **REF** | **ALT** |
| --- | --- | --- | --- | --- |
| PEKT02000001 | 282386 | B9J08_000131 | C | A |
| PEKT02000001 | 560007 | B9J08_000270 | T | G |
| PEKT02000001 | 610303 | B9J08_000297 | A | G |
| PEKT02000001 | 639957 | B9J08_000315 | A | G |
| PEKT02000001 | 683679 | B9J08_000336 | G | C |
| PEKT02000001 | 683682 | B9J08_000336 | T | G |
| PEKT02000001 | 683691 | B9J08_000336 | G | A |
| PEKT02000001 | 690603 | B9J08_000338 | A | C |
| PEKT02000001 | 704346 | B9J08_000345 | C | T |
| PEKT02000001 | 704362 | B9J08_000345 | T | G |
| PEKT02000001 | 704370 | B9J08_000345 | A | C |
| PEKT02000001 | 860319 | B9J08_000420 | A | G |
| PEKT02000001 | 860320 | B9J08_000420 | G | C |
| PEKT02000001 | 920410 | B9J08_000440 | T | G |
| PEKT02000001 | 921000 | B9J08_000440 | T | A |
| PEKT02000002 | 122994 | B9J08_000562 | T | C |
| PEKT02000002 | 227889 | B9J08_000611 | G | T |
| PEKT02000002 | 232332 | B9J08_000612 | T | G |
| PEKT02000002 | 355970 | B9J08_000675 | A | G |
| PEKT02000002 | 392101 | B9J08_000690 | G | A |
| PEKT02000002 | 448740 | B9J08_000714 | T | C |
| PEKT02000002 | 455561 | B9J08_000717 | T | A |
| PEKT02000002 | 489011 | B9J08_000733 | T | A |
| PEKT02000002 | 551157 | B9J08_000760 | A | C |
| PEKT02000002 | 897883 | B9J08_000921 | A | C |
| PEKT02000002 | 897884 | B9J08_000921 | T | C |
| PEKT02000002 | 897888 | B9J08_000921 | T | A |
| PEKT02000002 | 897893 | B9J08_000921 | C | T |
| PEKT02000002 | 897902 | B9J08_000921 | C | T |
| PEKT02000002 | 897918 | B9J08_000921 | C | T |
| PEKT02000002 | 926066 | B9J08_000929 | T | G |
| PEKT02000002 | 927719 | B9J08_000929 | T | G |
| PEKT02000002 | 991428 | B9J08_000958 | G | A |
| PEKT02000003 | 47310 | B9J08_001082 | C | A |
| PEKT02000003 | 194135 | B9J08_001155 | G | T |
| PEKT02000003 | 229610 | B9J08_001170 | A | C |
| PEKT02000003 | 281298 | B9J08_001195 | A | G |
| PEKT02000003 | 316781 | B9J08_001214 | T | A |
| PEKT02000003 | 362184 | B9J08_001235 | T | G |
| PEKT02000003 | 420359 | B9J08_001265 | C | A |
| PEKT02000003 | 494343 | B9J08_001296 | T | G |
| PEKT02000003 | 533174 | B9J08_001307 | T | G |
| PEKT02000003 | 638068 | B9J08_001359 | A | G |
| PEKT02000003 | 646318 | B9J08_001363 | C | G |
| PEKT02000003 | 666614 | B9J08_001370 | A | T |
| PEKT02000003 | 726826 | B9J08_001397 | T | G |
| PEKT02000003 | 738557 | B9J08_001404 | C | T |
| PEKT02000003 | 825765 | B9J08_001443 | C | T |
| PEKT02000003 | 917590 | B9J08_001479 | G | C |
| PEKT02000004 | 114850 | B9J08_001578 | C | T |
| PEKT02000004 | 251951 | B9J08_001647 | A | C |
| PEKT02000004 | 601993 | B9J08_001810 | C | G |
| PEKT02000004 | 734124 | B9J08_001882 | C | G |
| PEKT02000004 | 734126 | B9J08_001882 | C | T |
| PEKT02000004 | 734127 | B9J08_001882 | T | C |
| PEKT02000004 | 734129 | B9J08_001882 | T | C |
| PEKT02000005 | 31862 | B9J08_001955 | T | G |
| PEKT02000005 | 97587 | B9J08_001987 | T | G |
| PEKT02000005 | 105641 | B9J08_001991 | A | C |
| PEKT02000005 | 317956 | B9J08_002092 | C | G |
| PEKT02000005 | 317958 | B9J08_002092 | T | C |
| PEKT02000005 | 413870 | B9J08_002136 | T | G |
| PEKT02000005 | 533737 | B9J08_002188 | T | G |
| PEKT02000005 | 636353 | B9J08_002235 | A | G |
| PEKT02000006 | 21915 | B9J08_002245 | C | T |
| PEKT02000006 | 22263 | B9J08_002245 | A | T |
| PEKT02000006 | 124513 | B9J08_002288 | A | C |
| PEKT02000006 | 187702 | B9J08_002313 | A | C |
| PEKT02000006 | 187825 | B9J08_002313 | A | T |
| PEKT02000006 | 573599 | B9J08_002487 | A | T |
| PEKT02000006 | 613144 | B9J08_002504 | T | G |
| PEKT02000006 | 619454 | B9J08_002507 | A | C |
| PEKT02000006 | 710441 | B9J08_002547 | A | T |
| PEKT02000007 | 128130 | B9J08_002647 | G | A |
| PEKT02000007 | 231206 | B9J08_002682 | A | G |
| PEKT02000007 | 329125 | B9J08_002735 | C | G |
| PEKT02000007 | 485263 | B9J08_002816 | T | G |
| PEKT02000007 | 614617 | B9J08_002871 | C | G |
| PEKT02000007 | 763660 | B9J08_002949 | G | C |
| PEKT02000007 | 1057427 | B9J08_003111 | A | C |
| PEKT02000007 | 1471164 | B9J08_003310 | C | T |
| PEKT02000007 | 1690751 | B9J08_003417 | A | C |
| PEKT02000007 | 1823772 | B9J08_003473 | G | C |
| PEKT02000007 | 1850770 | B9J08_003489 | T | A |
| PEKT02000007 | 1893524 | B9J08_003509 | T | C |
| PEKT02000007 | 1943524 | B9J08_003539 | A | T |
| PEKT02000007 | 1995598 | B9J08_003564 | T | C |
| PEKT02000007 | 2007198 | B9J08_003571 | A | G |
| PEKT02000007 | 2214997 | B9J08_003661 | A | G |
| PEKT02000007 | 2252372 | B9J08_003683 | C | G |
| PEKT02000007 | 2252373 | B9J08_003683 | A | C |
| PEKT02000007 | 2270847 | B9J08_003694 | T | G |
| PEKT02000007 | 2353547 | B9J08_003729 | G | C |
| PEKT02000007 | 2412923 | B9J08_003761 | T | G |
| PEKT02000007 | 2462216 | B9J08_003780 | A | G |
| PEKT02000007 | 2483605 | B9J08_003788 | T | G |
| PEKT02000007 | 2578037 | B9J08_003829 | C | G |
| PEKT02000007 | 2719601 | B9J08_003891 | T | G |
| PEKT02000007 | 2731321 | B9J08_003896 | G | A |
| PEKT02000007 | 2752122 | B9J08_003906 | C | A |
| PEKT02000007 | 2752233 | B9J08_003906 | T | G |
| PEKT02000007 | 2754362 | B9J08_003907 | T | A |
| PEKT02000007 | 2823030 | B9J08_003936 | A | T |
| PEKT02000007 | 2931918 | B9J08_003986 | C | G |
| PEKT02000007 | 3009777 | B9J08_004023 | A | T |
| PEKT02000007 | 3026051 | B9J08_004027 | C | A |
| PEKT02000007 | 3049345 | B9J08_004036 | G | A |
| PEKT02000007 | 3049947 | B9J08_004037 | T | A |
| PEKT02000007 | 3081123 | B9J08_004053 | A | T |
| PEKT02000007 | 3081124 | B9J08_004053 | C | G |
| PEKT02000008 | 130012 | B9J08_004140 | G | A |
| PEKT02000008 | 268038 | B9J08_004193 | A | T |
| PEKT02000008 | 418422 | B9J08_004262 | A | C |
| PEKT02000008 | 481469 | B9J08_004290 | A | T |
| PEKT02000008 | 555186 | B9J08_004321 | T | A |
| PEKT02000008 | 555187 | B9J08_004321 | C | T |
| PEKT02000008 | 589823 | B9J08_004337 | A | C |
| PEKT02000008 | 622710 | B9J08_004349 | C | G |
| PEKT02000008 | 767238 | B9J08_004417 | G | A |
| PEKT02000008 | 771200 | B9J08_004419 | G | A |
| PEKT02000008 | 859650 | B9J08_004453 | T | C |
| PEKT02000008 | 860672 | B9J08_004453 | C | T |
| PEKT02000008 | 896291 | B9J08_004469 | G | A |
| PEKT02000009 | 78087 | B9J08_004498 | T | A |
| PEKT02000009 | 318567 | B9J08_004597 | A | G |
| PEKT02000009 | 383775 | B9J08_004627 | T | G |
| PEKT02000009 | 654838 | B9J08_004755 | A | C |
| PEKT02000009 | 752231 | B9J08_004790 | A | C |
| PEKT02000009 | 877590 | B9J08_004851 | T | C |
| PEKT02000009 | 944214 | B9J08_004877 | C | G |
| PEKT02000010 | 33333 | B9J08_004907 | G | A |
| PEKT02000010 | 50767 | B9J08_004913 | A | C |
| PEKT02000010 | 146276 | B9J08_004959 | T | G |
| PEKT02000010 | 174703 | B9J08_004971 | T | C |
| PEKT02000010 | 486030 | B9J08_005119 | T | C |
| PEKT02000010 | 513047 | B9J08_005131 | T | A |
| PEKT02000010 | 588985 | B9J08_005173 | G | T |
| PEKT02000010 | 624828 | B9J08_005191 | A | C |
| PEKT02000010 | 814666 | B9J08_005272 | C | T |
| PEKT02000010 | 816912 | B9J08_005273 | G | A |
| PEKT02000010 | 914390 | B9J08_005319 | A | C |
| PEKT02000010 | 955034 | B9J08_005337 | A | C |
| PEKT02000010 | 960587 | B9J08_005340 | C | G |
| PEKT02000010 | 1001210 | B9J08_005357 | T | G |
| PEKT02000010 | 1089035 | B9J08_005394 | T | G |
| PEKT02000010 | 1301342 | B9J08_005490 | G | C |
| PEKT02000010 | 1347020 | B9J08_005512 | A | T |
| PEKT02000012 | 2621 | B9J08_005545 | T | A |
| PEKT02000012 | 11113 | B9J08_005550 | T | A |
| PEKT02000014 | 3174 | B9J08_005579 | G | C |

**Table S6:** List Non-synonymous mutations present exclusively in A1

| **Chromosome** | **Positon** | **Gene ID** | **REF** | **ALT** |
| --- | --- | --- | --- | --- |
| PEKT02000001 | 526073 | B9J08_000250 | G | C |
| PEKT02000001 | 560044 | B9J08_000270 | A | T |
| PEKT02000002 | 283422 | B9J08_000639 | CTC | TTT |
| PEKT02000002 | 890952 | B9J08_000917 | GAAA | GAAG |
| PEKT02000003 | 193630 | B9J08_001155 | C | T |
| PEKT02000003 | 797353 | B9J08_001429 | TTC | TTT |
| PEKT02000004 | 395335 | B9J08_001712 | T | G |
| PEKT02000006 | 459602 | B9J08_002443 | G | T |
| PEKT02000006 | 699424 | B9J08_002544 | TGCCAGGAC | TGCCAGCGC |
| PEKT02000007 | 1535088 | B9J08_003347 | CAAAA | CATCA |
| PEKT02000007 | 2416354 | B9J08_003764 | CAAG | CAAA |
| PEKT02000007 | 2618295 | B9J08_003845 | A | G |
| PEKT02000007 | 3128406 | B9J08_004082 | C | A |
| PEKT02000007 | 3168758 | B9J08_004097 | A | T |
| PEKT02000007 | 3193627 | B9J08_004102 | A | C |
| PEKT02000008 | 388977 | B9J08_004249 | G | C |
| PEKT02000008 | 389569 |  | T | A |
| PEKT02000009 | 328635 | B9J08_004602 | A | T |
| PEKT02000010 | 588390 | B9J08_005173 | GAAC | AACA |
| PEKT02000012 | 10652 | B9J08_005550 | C | G |

**Table S7:** List Non-synonymous mutations present exclusively in A2

| **Chromosome** | **Positon** | **Gene ID** | **REF** | **ALT** |
| --- | --- | --- | --- | --- |
| PEKT02000001 | 526073 | B9J08_000250 | G | C |
| PEKT02000001 | 560044 | B9J08_000270 | A | T |
| PEKT02000002 | 283422 | B9J08_000639 | CTC | TTT |
| PEKT02000002 | 433854 | B9J08_000706 | AGTGC | AGTGG |
| PEKT02000003 | 851422 | B9J08_001455 | A | C |
| PEKT02000004 | 395335 | B9J08_001712 | T | G |
| PEKT02000006 | 21886 | B9J08_002245 | C | A |
| PEKT02000006 | 459602 | B9J08_002443 | G | T |
| PEKT02000006 | 699407 | B9J08_002544 | GCCAGGG | TGCTGCA |
| PEKT02000007 | 1535088 | B9J08_003347 | CAAAA | CATCA |
| PEKT02000007 | 1850769 | B9J08_003489 | ATCC | AACC |
| PEKT02000007 | 2416354 | B9J08_003764 | CAAG | CAAA |
| PEKT02000007 | 2618295 | B9J08_003845 | A | G |
| PEKT02000007 | 2941309 | B9J08_003990 | TGCTG | CGCTA |
| PEKT02000007 | 3128794 | B9J08_004082 | G | T |
| PEKT02000008 | 42005 | B9J08_004112 | G | C |
| PEKT02000008 | 389569 | B9J08_004249 | T | A |
| PEKT02000009 | 328635 | B9J08_004602 | A | T |
| PEKT02000010 | 588390 | B9J08_005173 | GAAC | AACA |
| PEKT02000012 | 10652 | B9J08_005550 | CTTG | GTTA |

**Table S8:** List Non-synonymous mutations present exclusively in A3

| **Chromosome** | **Positon** | **Gene ID** | **REF** | **ALT** |
| --- | --- | --- | --- | --- |
| PEKT02000001 | 526073 | B9J08_000250 | G | C |
| PEKT02000001 | 560044 | B9J08_000270 | A | T |
| PEKT02000001 | 684960 | B9J08_000336 | ACTG | GCTA |
| PEKT02000001 | 699725 | B9J08_000341 | AC | CG |
| PEKT02000002 | 353041 | B9J08_000675 | T | G |
| PEKT02000003 | 646078 | B9J08_001363 | AGAC | GGAG |
| PEKT02000003 | 646087 | B9J08_001363 | AGAC | GGAG |
| PEKT02000003 | 866909 | B9J08_001458 | AGT | GAA |
| PEKT02000006 | 21886 | B9J08_002245 | C | A |
| PEKT02000006 | 459602 | B9J08_002443 | G | T |
| PEKT02000006 | 699407 | B9J08_002544 | GCCAGGGC | TGCTGCAC |
| PEKT02000006 | 775394 | B9J08_002584 | G | C |
| PEKT02000007 | 1850769 | B9J08_003489 | ATCC | AACC |
| PEKT02000007 | 2618295 | B9J08_003845 | A | G |
| PEKT02000007 | 2941309 | B9J08_003990 | TGCTG | CGCTA |
| PEKT02000007 | 3128406 | B9J08_004082 | C | A |
| PEKT02000007 | 3128794 | B9J08_004082 | G | T |
| PEKT02000007 | 3130873 | B9J08_004083 | C | A |
| PEKT02000007 | 3168758 | B9J08_004097 | A | T |
| PEKT02000008 | 388977 | B9J08_004249 | G | C |
| PEKT02000009 | 328635 | B9J08_004602 | A | T |
| PEKT02000010 | 7854 | B9J08_004897 | G | A |
| PEKT02000010 | 588390 | B9J08_005173 | GAAC | AACA |
| PEKT02000012 | 10652 | B9J08_005550 | CTTG | GTTA |
| PEKT02000013 | 24127 | B9J08_005575 | C | A |
